# Synchronized swarming: Harmonic convergence and acoustic mating dynamics in the malaria mosquito *Anopheles gambiae*

**DOI:** 10.1101/2021.07.03.451017

**Authors:** Stefano S. Garcia Castillo, Kevin S. Pritts, Raksha S. Krishnan, Laura C. Harrington, Garrett P. League

## Abstract

The mosquito *Anopheles gambiae* is a major African malaria vector, transmitting parasites responsible for significant mortality and disease burden. Malaria declines have stagnated recently due to widespread insecticide resistance among vector populations. Flight acoustics are essential to mosquito mating biology and represent promising alternative targets for mosquito control. However, mosquito swarm acoustics data are limited. Here, for the first time, we present detailed analyses of free-flying male and female *An. gambiae* flight tones and their harmonization (harmonic convergence) over a complete swarm sequence. Audio analysis of single-sex swarms showed elevated male or female flight tone frequencies and amplitudes during swarming flight with gradual declines to pre-swarm levels over an approximately 35-min period. Analysis of mixed-sex swarms revealed additional increases in flight tone frequencies and amplitudes due to mating activity. Data from mixed-sex swarms suggest harmonic convergence during swarming enhances the efficiency of female detection by synchronizing male and female baseline swarm tones. Further, data from experiments using female swarm tone playbacks to males indicate that harmonic convergence during mating interactions coordinates male scramble competition by acoustically masking mating couple flight tones. These findings advance our knowledge of mosquito swarm acoustics, providing vital information for reproductive control strategies.

## Introduction

Eradication of mosquito-borne illness like malaria remain a top global public health priority. In 2019 there were an estimated 229 million malaria cases globally, the vast majority of which occurred in Africa (94%), and 409,000 malaria deaths, mainly involving children under the age of five years (67%)^1^. Currently, the most widely adopted malaria control efforts employ insecticides to target the mosquito vector by means of long-lasting insecticide-treated bed nets and indoor residual spraying. Although these tactics have led to marked declines in malaria from the year 2000 to approximately 2015, these declines have plateaued due to the spread of insecticide resistance in malaria vector mosquitoes^1^. Novel vector and disease control strategies are thus urgently needed to overcome the stagnating progress of these chemical-based control methods.

Anopheline mosquitoes are solely responsible for human malaria transmission. Among these mosquitoes, *Anopheles gambiae* represents one of the most efficient malaria vectors in sub-Saharan Africa. As such, significant effort had been devoted to studying *An. gambiae* biology and behavior as a means of developing effective disease control strategies. Reproductive control methods have gained traction in recent years because of their capacity to interrupt mating to reduce or eliminate vector populations. Mating in *An. gambiae* occurs in flight in swarms composed of tens, hundreds, or even thousands of individuals, the vast majority of which are male^2–6^. Swarms occur primarily at dusk^4, 7^ and near human habitations, to increase female encounter rates^6^. Swarming starts with a few males flying in a “dancing” flight pattern^8^ followed by a gradual influx of additional swarming males^9^. Upon formation of a swarm, males locate potential female mates using differences between male and female flight tones^10^. Once paired, females drop and/or fly out of the swarm with the male attached in copula^2, 4, 6^ while males transfer semen and a mating plug into the female in under 20 sec^11^ to prevent further copulations^12, 13^.

Because flight tones are essential to mosquito mating communication, several recent studies have successfully utilized mosquito flight tones for trapping and monitoring purposes^14–24^. However, detailed information on swarm acoustics is lacking due to the technical challenges associated with recording and analyzing complex group mating sounds.

Previous studies of individual mosquito mating pairs have contributed substantially to our understanding of the dynamic interplay of male and female acoustics during mating interactions. During courtship flight that immediately precedes mating, a male and female will often actively match their wing beat frequencies at their nearest shared harmonic in a phenomenon called harmonic convergence^25–30^. Harmonic convergence has been documented in numerous mosquito species^26, 28, 29, 31^ and is essential in sex recognition^29, 32–35^, species recognition^31–35^, and even reproductive isolation and assortative mating^31–34, 36, 37^ (although this appears not to be the case for some closely related species like *An. gambiae* and *An. coluzzii*^38, 39^). As harmonic convergence also correlates strongly with mating success^25, 40, 41^, studies on sexual selection and mate preference suggest that convergence may serve as a signal of some^26, 42^, but not all^43^ aspects of mosquito fitness. Despite these discoveries, much of the previous work on harmonic convergence has examined tethered, rather than free-flying males and females. Although tethered flight analysis offers an experimentally tractable means of dissecting the mechanics and biological significance of this behavior, tethering restricts natural flight patterns and alters flight tones^10, 21, 44–48^. Analysis on flight tone acoustics and harmonic convergence over the course of a complete swarm sequence would therefore provide valuable insights and context for these phenomena, as studies examining convergence in free-flight are limited^10, 39, 47, 49^.

In the current study, we examine free-flight acoustics and harmonic convergence across an entire *An. gambiae* swarm sequence in high-resolution detail in both single and mixed-sex swarms. Importantly, we link swarm acoustics to their underlying mating behaviors, showing how the two are biologically interdependent. Surprisingly, we found that harmonic convergence and harmonic frequency differences of less than 50 Hz not occur not only between male and female swarm tones during mating attempts but also between collective male and female baseline swarm tones. Furthermore, although female baseline swarm tones are highly attractive to males, female tones during mating attempts, which are higher in frequency and amplitude, are almost completely unattractive. Together, our data suggest harmonic convergence serves as a mechanism for coordinating *An. gambiae* swarm mating by enhancing male detection of available females and acoustically masking mating pairs. These findings advance our understanding of mosquito swarm mating acoustics and provide additional supporting evidence for the utility of flight tones in reproductive control strategies.

## Methods

### Mosquito rearing and maintenance

*Anopheles gambiae* (G3) larvae and adults were reared and maintained in an incubator (Powers Scientific Inc., Pipersville, PA, USA) at 25.92 ± 1.17 °C and 67.44 ± 4.18 % relative humidity (RH). Similar to previous *An. gambiae* laboratory swarm studies^4, 9, 50^, the incubator was programmed with a 12 h light: 12 h dark photoperiod (two 17-watt fluorescent tube bulbs), including 1 hr of crepuscular lighting (one 5.5-watt LED bulb) at dawn and dusk. Because flight tone frequencies can vary by adult size^21, 24, 26, 32^, rearing methods were designed to obtain field-sized adult mosquitoes^6, 51–54^ of equivalent sizes between trials. Male and female wing lengths from a subset of individuals from each experimental group were measured as a proxy for overall size as previously described^55^ (Supplementary Fig. S1; Supplementary Table S1). For all experiments, 250 L1 larvae were transferred into 30 cm x 23 cm x 8 cm plastic trays containing 1 L of distilled water. Larvae were provided 52 mg of ground Cichlid fish food (Hikari Cichlid Gold, Hayward, CA, USA) per day from day one through four of development and 150 mg of food per day from day five of development until pupation (resulting in approximately 0.2 mg of fish food per larva per day)^56^. Pupae were sexed via examination of the terminalia and placed separately by sex and age into 2 L wax-lined cardboard buckets covered with mosquito netting. Adults were provided with a cotton pad soaked in a 10% sucrose solution. Experiments were conducted on adults at 4–7 days post-eclosion, the ideal swarming and mating age range for sexually mature males^53^.

### Swarm cage setup and audio recording and analysis

Swarm recordings were conducted under tightly-controlled, artificially-inducible swarming conditions to recapitulate natural mating behavior as in previous laboratory studies^9, 50, 57^. To control for potential effects of temperature on flight tone frequencies and swarming activity^10, 21, 24, 58^, all swarm recordings took place at 26.35 ± 2.18 °C and 63.80 ± 10.94 % RH at the beginning of the scotophase (i.e., dark phase following dusk) when swarm mating activity occurs in nature^4, 7^. Recordings took place in a 14 cm x 14 cm x 14 cm steel-framed cage containing four mosquito net side panels, a steel perimeter lid with a mosquito net cover, and a cardstock paper base with a 4 cm x 4 cm black paper swarm marker placed at its center as in previous setups^4, 9, 50^ (Supplementary Fig. S2). A small incision was made in the netting to insert an omnidirectional microphone (FG-23329-C05; Knowles Electronics, Itasca, IL, USA) inside the cage above the middle of the right border of the swarm marker. The microphone was attached to a custom amplifier^59^, which was connected via USB port to a nearby computer. Ambient recording room lighting (two 35-watt LED tube bulbs) was used to simulate dusk for the first 15 min of the recording. Dusk lights were then turned off for the remaining 45 min of the recording to mimic sunset and the initial period of night, when male antennal fibrillae are erect to optimize detection of females^4, 11, 31^ and circadian flight activity peaks^58, 60, 61^. To observe and record mating activity while minimizing light interference for the mosquitoes, the swarm cage was backlit throughout the recording with a red-light lamp^62^ covered with a white paper diffuser. A white paper-lined recording booth was also placed around the swarm cage to optimize illumination. This booth was itself placed within a wooden, acoustic-foam-lined mosquito resting box to minimize external noise interference. Videos were taken using a FITFORT 4K Action Camera (Dreamlink E-Commerce Co., Shenzhen, GD, CN) to visualize and verify swarm mating parameters recorded manually during audio recordings.

Mosquitoes acclimated in the recording cage for 5 min prior to commencement of recordings. Audio and video recordings encompassed the entire 1-hr-long swarm sequence, including a pre-swarm period (0–15 min) in dusk conditions, and a peak-swarm (15–30 min), waning-swarm (30–45 min), and late-swarm period (45–60 min) in dark conditions. Recordings were obtained using Audacity software version 2.1.3^63^. Spectrograms from audio obtained in Audacity were analyzed using Raven Pro 1.5 software^64^ (Bioacoustics Research Program, Cornell Laboratory of Ornithology, Ithaca, NY, USA) to determine male (M1) and female (F1) fundamental flight tone frequencies (Hz) and amplitudes (dB). To reduce noise and enhance swarm tone signals, a 10 sec silent audio recording was acquired prior to addition of mosquitoes into the recording cage to perform a background noise subtraction using Audacity’s "Noise Reduction" effect.

### Single-sex swarm experiments

For single-sex swarm experiments, swarms consisting of either 40 male or 40 female 4– 7-day-old mosquitoes were recorded. A total of six single-sex swarms were recorded, with three trials conducted with males and three trials conducted with females. For each 1-hr recording, collective male or individual female fundamental flight tone frequencies and amplitudes were sampled from audio spectrograms every 60 sec and data were binned by swarm period (180 male and 180 female samples total across all trials).

### Mixed-sex swarm experiments

For mixed-sex swarm experiments, swarms consisting of 30 male and 10 female (3:1 male:female swarm sex ratio^65^) age-matched mosquitoes were recorded. A total of 12 mixed-sex swarms were recorded, with three trials conducted for each of the four mosquito age groups (4, 5, 6, and 7-day-olds). The timing of each mating attempt (male-female mating interaction that led to the couple dropping out of the swarm) and copula (male and female genitalia connected either in flight or on a cage surface for >10 s^11^; Supplementary Fig. S2) was recorded simultaneously using an Android (version 7.1.1) Stopwatch app (Android, Palo Alto, CA, USA) and the Stopwatch and Timer app (Jupiter Apps, Dover, Kent, UK). Immediately after each recording, females were dissected on an Olympus SZX10 stereo microscope (Olympus Corp., Shinjuku City, Tokyo, JP) to determine mating status. Insemination status was determined by the presence or absence of sperm in the spermatheca and successful mating plug transfer was determined by the presence or absence of an internal mating plug (Supplementary Fig. S3). Dissections were imaged under oblique, darkfield, and brightfield illumination, as well as fluorescence illumination to visualize autofluorescent mating plugs using a GFP filter set (SZX-MGFPA, Olympus) and an X-Cite 120 Fluorescence Illumination System (EXFO, Quebec City, Quebec, CA) on maximum excitation setting. Images of dissections (and the swarm cage setup above) were obtained using a Sony IMX214 Exmor R CMOS 13-megapixel camera with optical image stabilization and f/2.0 aperture (Sony Corp., Minato City, Tokyo, JP).

Male and female flight tone frequencies and amplitudes were sampled and binned as with single-sex swarm recordings. Additionally, we checked for harmonic convergence, defined as flight tone harmonic differences of <5 Hz^25^, in male-female acoustic mating interactions (Supplementary Fig. S4) every 5 min from 15–60 min (ten interactions per trial for twelve mixed-sex swarm trials for 120 total sampled interactions across all trials). If more than one mating interaction or mating interaction segment occurred at a time point, only the first segment of the interaction closest to the sampling time point was analyzed (Supplementary Fig. S4). To be considered a mating interaction tone, male tones had to display each of the following three criteria: 1) a ≥ 25 Hz increase in frequency relative to background swarming male frequencies; 2) a ≥10 dB increase in amplitude relative to background male tones; and 3) rapid frequency modulation, a distinctive behavioral feature of male mosquito mating interaction flight^10, 39, 40, 47, 48, 66^. Harmonic overtones and flight tone harmonic differences were estimated based on sampling from fundamental flight tone frequencies as previously described^67^. However, the <5 Hz criterion for harmonic convergence was based on experiments with tethered mosquitoes^27^ and free-flight interactions afford briefer periods (often only ≈1–2 sec^10, 35, 39, 47^) in which to assay for harmonic convergence^49^. Furthermore, we found that flight tone harmonic differences greater than 50 Hz were sufficient to distinguish courting couple tones from swarming male and female tones (see Results, Figs. 6, 7). Thus, in addition to retaining the previously established <5 Hz harmonic difference criterion, we also included difference outcomes that were <50 but >20 Hz, <20 but >10 Hz, and <10 but >5 Hz in our analyses. Collectively, these four outcome categories (i.e., <50, <20, <10, and <5 Hz) are referred to as harmonic frequency difference outcomes (or some variation of this).

We assayed male and female tones for harmonic convergence during both the early (“Early interaction” phase) and late (“Late interaction” phase) phases of acoustic mating interactions (Supplementary Fig. S4). Flight tones were assayed at three time points near the beginning, middle, and end of each interaction phase (120 before and 120 during phases for 240 total phases across all trials). Harmonic convergence was assayed at four harmonic ratios typically associated with convergence^26, 27, 40, 67^: 1) the male first and female second harmonics (M1:F2); 2) the male second and female third harmonics (M2:F3); 3) the male third and female fourth harmonics (M3:F4); and 4) the male third and female fifth harmonics (M3:F5). For each of the two interaction phases (i.e., early and late interaction) harmonic difference outcomes occurred in three possible scenarios: a harmonic difference occurred both early and late in an interaction (scenario 1), only early in an interaction (scenario 2), or only late in an interaction (scenario 3). Interacting mosquitoes that did not achieve one of the four harmonic difference categories in either the early or late mating interaction phases were placed in a separate “>50 Hz” outcome scenario (Supplementary Data S1). Time points where no interacting mosquitoes were present (three instances, all at 60 min) were also placed in the “>50 Hz” category. Outcome scenarios were categorized by harmonic differences as well as the harmonic ratio, age, and swarm period in which these differences occurred.

### Single-sex swarm experiments with audio playback

Artificial swarm tone frequency and amplitude parameters were determined by sampling male and female tones from a random subset of single- and mixed-sex swarm trials (for details, see Supplementary File S1). For audio playback of artificial female swarm tones to single-sex swarms consisting of 40 males (three trials), a 1-hr artificial female swarm audio playback was created using Audacity software’s Generate Tone and Add New Track functions to generate fundamental tones (F1) and harmonic overtone stacking (F2–F5). Artificial female swarm audio consisted of an initial 15 min of silence followed by 45 min of alternating 5 sec periods of either female baseline swarm tone (550 Hz) or female mating interaction tone (600 Hz) frequencies (270 baseline swarm tone periods and 270 mating interaction tone periods). Artificial swarm audio was played from an Apple iPod MA662G/B Headphone (Apple Inc., Cupertino, CA, USA) speaker featuring a 20–20,000 Hz frequency response placed 2 cm from the microphone (Supplementary Fig. S2).

Male fundamental flight tone frequencies and amplitudes were sampled from female artificial playback swarm spectrograms every 60 sec for the first 15 min of silence (15 samples) and every 60 sec during both a baseline swarm tone and mating interaction tone female playback period for the next 45 min (90 samples for 105 frequency and amplitude samples total per trial and 315 samples total across all trials). If present, we sampled tones when males engaged in a mating interaction (n=34 instances; see above for criteria); if interacting males were not present, we instead sampled baseline male swarm tones from the midpoint of each 5 sec baseline swarm tone and mating interaction tone female playback period at that time point. For comparisons between interacting and non-interacting male flight tone parameters, male interaction tones, which occurred almost exclusively during female baseline swarm tone playback periods, were compared to the nearest non-interacting male tone obtained during a female baseline swarm tone period. We also tested for the presence or absence of male mating interactions at all 540 artificial female swarm audio playback periods (270 baseline swarm tone periods and 270 mating interaction tone periods) for a total of 1620 samples over three trials.

Further, we assayed harmonic differences for all mating interactions detected during the 45 min alternating female tone playback period at M2:F3, the most common ratio for harmonic differences from the mixed-sex swarms trails (see Results, Figs. 2, 5). Similar to the mixed-sex swarms, for each mating interaction harmonic differences between background male swarm tones and either the female baseline swarm tone or mating interaction tone were assayed at three time points immediately before the interaction (“Before interaction” phase; see Results, Fig. 7). Then, harmonic frequency differences between interacting male tones and either the female baseline swarm tone or mating interaction tone were assayed at three time points during the interaction (“During interaction” phase; Fig. 7). For each of the four interaction phases (i.e., before interaction and female baseline swarm tone, before interaction and female mating interaction tone, during interaction and female baseline swarm tone and during interaction and female mating interaction tone), harmonic differences occurred in twelve possible outcome scenarios (see Results, Fig. 8).

For playback of artificial male swarm tones to single-sex swarms consisting of 40 females (three trials), a 1-hr artificial male swarm audio playback was created using the same method used to create artificial female swarm audio (Supplementary File S1). Artificial male swarm audio consisted of 15 min of silence followed by 45 min of alternating 5 sec periods of either male baseline swarm tone (850 Hz) or male mating interaction tone (900 Hz) frequencies (270 baseline swarm tone periods and 270 mating interaction tones periods). Female fundamental flight tone frequencies and amplitudes were sampled from male artificial playback swarm spectrograms every 60 sec for the first 15 min of silence (15 samples) and every 60 sec during both a baseline swarm tone and mating interaction tone male playback period for the next 45 min (90 samples for 105 frequency and amplitude samples total per trial and 315 samples total across all trials). Because female mosquitoes never attempted to mate with the speaker emitting the artificial male audio, we did not examine mating interaction acoustics or harmonic convergence in audio playback experiments to female swarms.

### Statistical analysis

Data were analyzed with R statistical software (Version 4.0.0) in RStudio software (Version 1.2.5042) with model packages "rstatix"^68^, “ggplot2”^69^, “ggpubr”^70^, "ComplexUpset"^71^, and "RVAideMemoire"^72^. For all analyses, p-values <0.05 were considered statistically significant. Significant differences were denoted using lowercase APA-style letter subscripts. For box and whisker plots throughout, boxes display the boundaries of the first (bottom) and third (top) quartiles, median lines, and individual data points (including outliers) and whiskers denote the minimum and maximum values. Numerical data are presented as average ± standard deviation (SD) throughout. Raw numerical data for all figures are included in Supplementary Data S1.

One-way repeated measures ANOVAs with a Tukey post-hoc test were used to test for differences in flight tone frequencies, flight tone amplitudes, mating attempts, and copulas between swarm periods. For tests involving independent variables, such as age and artificial audio playback period (baseline swarm tone or mating interaction tone), two-way repeated measures ANOVAs with a Tukey post-hoc test were used. A Fisher’s exact test was used to evaluate differences in the proportion of inseminated females or females with a mating plug between ages, the proportion of harmonic frequency differences between mating interaction phases and female audio playback periods, as well as the proportion of male mating interaction tones that met each of the four mating interaction criteria. To evaluate the proportion of harmonic frequency differences (and/or the harmonic ratios, ages, and swarm periods in which they occurred) within and between outcome scenarios, as well as the proportion of male mating interactions that occurred between female audio playback periods, a Chi-square goodness-of-fit test was performed using the same expected probability for all groups followed by a Benjamini-Hochberg (B-H) post-hoc test^73^ when appropriate. A paired t-test was used to compare male flight tone frequencies and amplitudes in the presence or absence of a mating interaction.

## Results

### Male and female single-sex swarm tones climax early in swarming, decrease gradually over time, and differ in degree of modulation

To track male and female swarm tone modulations across an entire swarm sequence in the absence of the opposite sex, we measured flight tone frequencies and amplitudes in single-sex swarms (Fig. 1). Male swarm tone frequencies (one-way repeated measures ANOVA, F(3,176)=21.29, p<0.0001) and amplitudes (ANOVA, F(3,176)=23.73, p<0.0001) differed significantly between swarm periods, climaxing immediately following sunset in the peak-swarm period (15–30 min) and returning to pre-swarm period (0–15 min) values in the waning-swarm (30–45 min) and late-swarm periods (45–60 min; Fig. 1A–D; Tukey post-hoc test; Supplementary Table S2).

**Figure 1.**
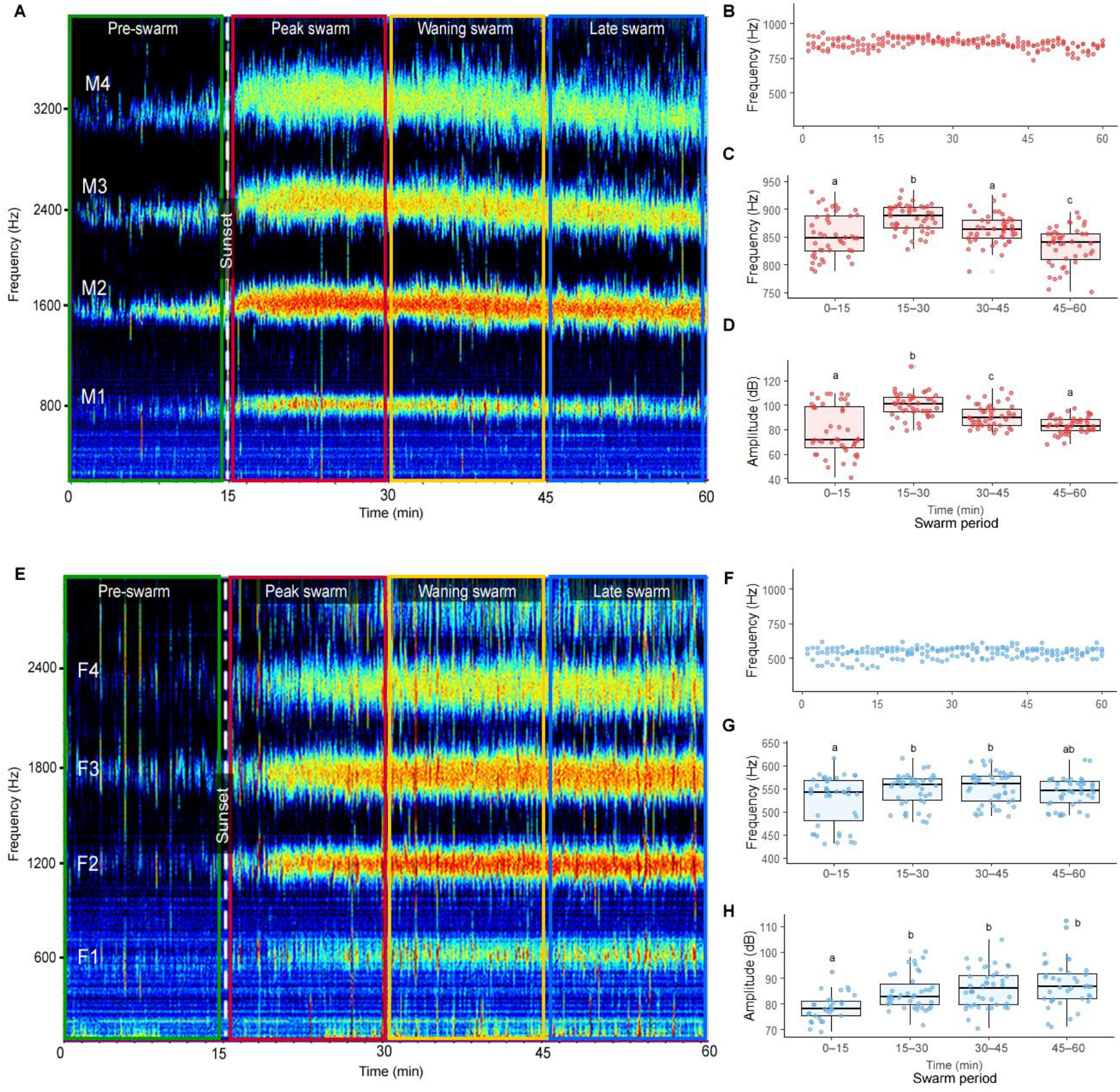
Male and female single-sex swarm tones vary by swarm period and sex. (**A**) Representative male single-sex swarm audio spectrogram. Male fundamental swarm tone frequencies (**B**, **C**) and amplitudes (**D**) analyzed by swarm period. (**E**) Representative female single-sex swarm audio spectrogram. Female fundamental swarm tone frequencies (**F**, **G**) and amplitudes (**H**) analyzed by swarm period. Lowercase letters above box and whisker plots denote Tukey post-hoc test p-values for comparisons between swarm periods. Abbreviations: M1, male fundamental flight tone harmonic; M2, male second harmonic; M3, male third harmonic; M4, male fourth harmonic; F1, female fundamental flight tone harmonic; F2, female second harmonic; F3, female third harmonic; F4, female fourth harmonic.

Female flight tone modulations over a swarm sequence were analogous to males, though less pronounced (Fig. 1E–H; Supplementary Table S2). Female swarm tone frequencies (one-way repeated measures ANOVA, F(3,174)=4.15, p<0.01) and amplitudes (ANOVA, F(3,146)=9.613, p<0.0001) differed between swarm periods (Tukey post-hoc test). Although elevated female peak-swarm frequencies partially returned to pre-swarm levels in the late-swarm period (Fig. 1G), elevated female swarm amplitudes did not (Fig. 1H; Tukey post-hoc test). Consistent with earlier studies in *Anopheles*^32, 36^ and other mosquito species^10, 28, 37, 74^, female swarm tone frequencies were approximately 300 Hz (i.e., ≈1.5x) lower than males flight tone frequencies (Supplementary Table S2). Together, these swarm recordings show that both male and female flight tone parameters vary substantially by swarm period and sex, even in the absence of the opposite sex.

### Male and female mixed-sex swarm flight tones display significant harmonic overlap, vary by swarm period and sex, and increase with mating activity and age

To evaluated male and female swarm acoustics simultaneously across a complete swarm sequence, we measured flight tone parameters in mixed-sex swarms (Fig. 2). Despite reduced activity in both sexes before sunset, we observed consistent early swarming by a small contingent of males (5 ± 3 individuals), which we refer to here as sentinel swarmers (Fig. 2A; Supplementary File S2). Sentinel swarming occurred primarily over the 10-min period prior to sunset and occurred at lower, more variable frequencies compared to peak and waning swarm tones. Male swarming occurred continuously after sunset, albeit with declining participation over time, reaching peak frequencies ≈3 min after sunset (swarm latency) and returning to pre-swarm frequencies ≈35 min after peaking (swarm duration; Supplementary File S2). Female swarming consisted of relatively discontinuous bursts of “offering flight”^4^, which were quickly interrupted by male mating attempts. Perhaps due to frequent increases in male and female flight tone frequencies and amplitudes during mating interactions (Fig. S4), male and female swarm tone parameters were elevated in mixed-sex swarms compared to single-sex swarms (Supplementary Table S2 and S3). Male and female mixed-sex swarm tone frequencies also displayed significant overlap, particularly during peak-swarming, at the male second and female third harmonics (M2:F3), which both occur at ≈1,800 Hz (Fig. 2A).

**Figure 2.**
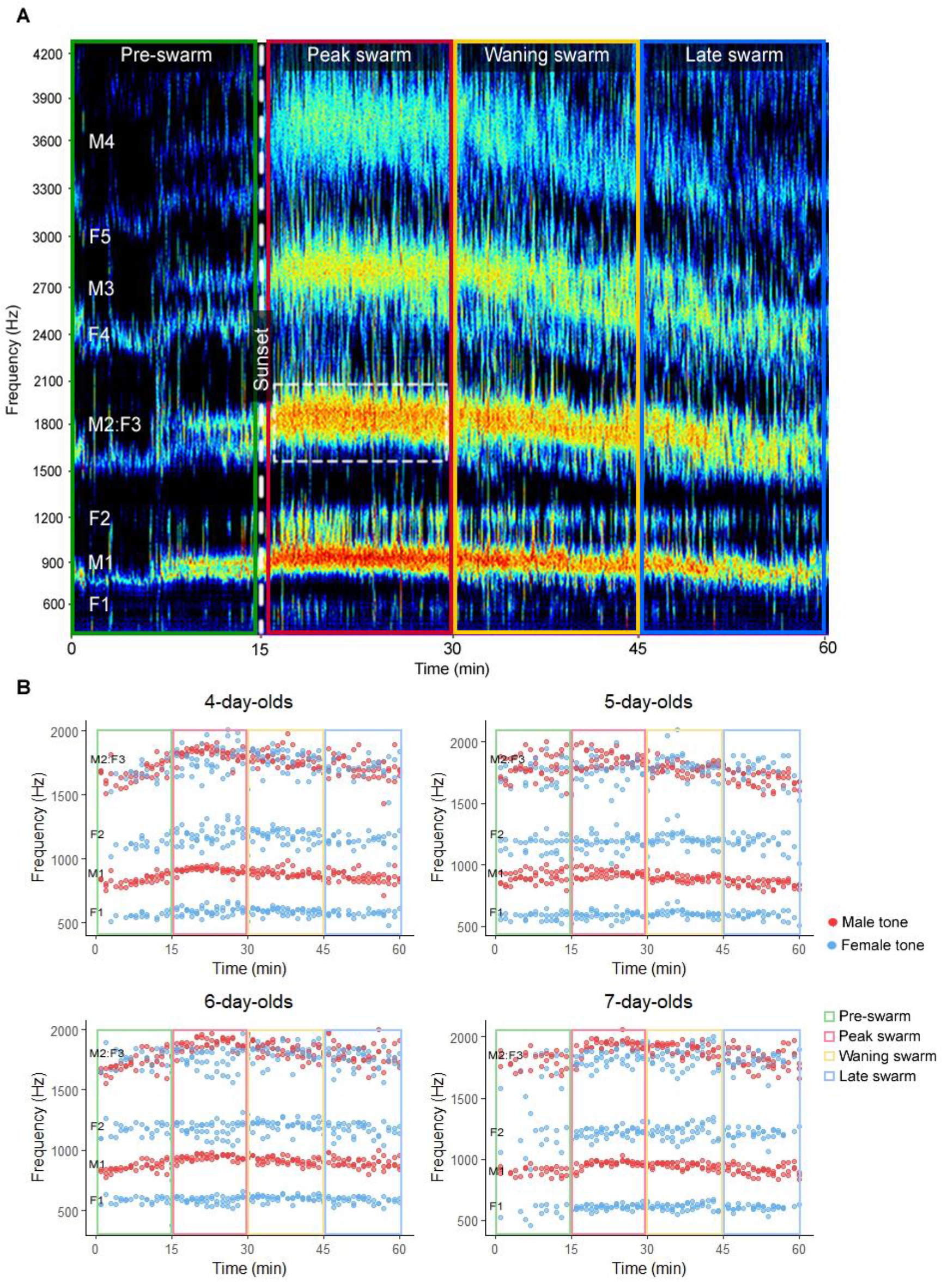
Male and female mixed-sex swarm tones display harmonic overlap. (**A**) Representative mixed-sex swarm audio spectrogram. White dashed box highlights a portion of the significant overlap between the male second and female third harmonics (M2:F3). (**B**) Male and female swarm tone frequencies displayed separately by age. Abbreviations: M1, male fundamental flight tone harmonic; M2, male second harmonic; M3, male third harmonic; M4, male fourth harmonic; F1, female fundamental flight tone harmonic; F2, female second harmonic; F3, female third harmonic; F4, female fourth harmonic; F5, female fifth harmonic.

Both male and to a lesser degree female mixed-sex swarm tone parameters varied significantly between swarm periods and adult ages, with frequencies climaxing during peak swarming, declining with time, and increasing with age from 4–7 days post-eclosion (Fig. 2B; Fig. 3; Supplementary Table S3 and S4; two-way repeated measures ANOVA; Tukey post-hoc test). Collectively, mixed-sex swarm recordings show that male and female swarm tones display substantial harmonic overlap and vary by swarm period, sex, mating activity, and age.

**Figure 3.**
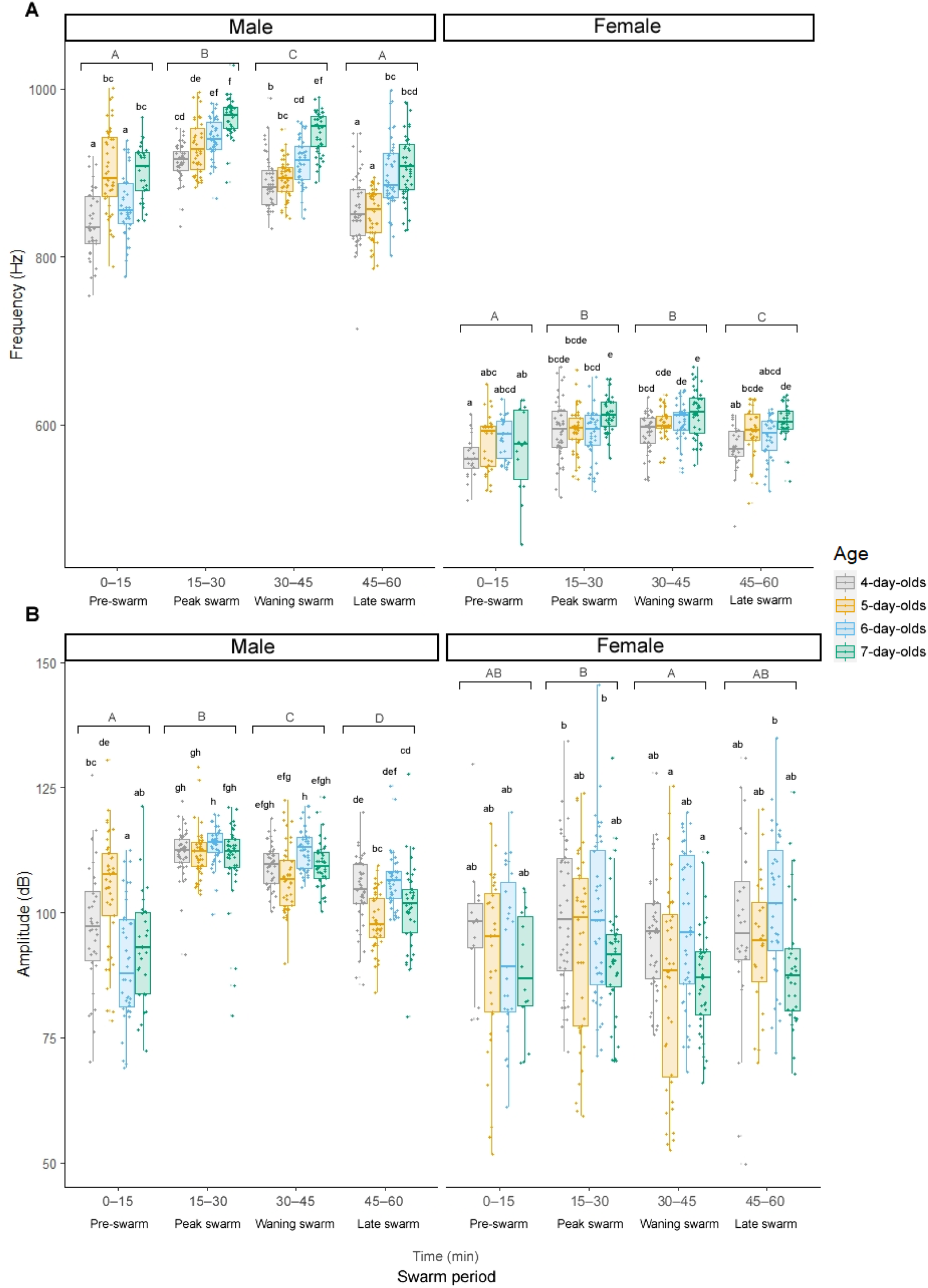
Male and female mixed-sex swarm tone parameters vary by swarm period, sex, and age. Male and female fundamental swarm tone frequencies (**A**) and amplitudes (**B**) by swarm period and age. Letters above box and whisker plots denote Tukey post-hoc test p-values for comparisons between swarm periods (uppercase) and ages (lowercase).

### Mating activity climaxes during the peak swarm period

Mixed-sex swarms displayed elevated swarm tone parameters during mating interactions. We therefore quantified mating attempts and copulas (Supplementary Fig. S2) across complete mixed-sex swarm sequences to compare trends in mating activity and swarm tone modulation (Fig. 4). As expected, mating attempts differed significantly by swarm period (two-way repeated measures ANOVA, F(3,128)=73.663, p<0.0001) and age (ANOVA, F(3,128)=2.901, p=0.038), increasing dramatically from the pre-swarm period (3.64 ± 9.23) to the peak swarm period (41.50 ± 21.00) and gradually declined in the waning (14.72 ± 8.99) and late (6.00 ± 5.12) swarm periods (Fig. 4A; Supplementary Table S5 and S6; Tukey post-hoc test). Mating copulas were almost entirely absent during pre-swarm period (0.06 ± 0.23), as seen in *Anopheles* swarms in nature^4^. As with mating attempts, copulation climaxed in the peak swarm period (2.44 ± 1.63) and declined gradually with time (waning swarm: 0.64 ±1.05; late swarm: 0.25 ± 0.55; Fig. 4B; Supplementary Table S5 and S6; two-way repeated measures ANOVA, F(3,128)=39.569, p<0.0001; Tukey post-hoc test). However, unlike mating attempts, copulation occurred at similar rates between ages (Supplementary Table S5 and S6; Tukey post-hoc test).

**Figure 4.**
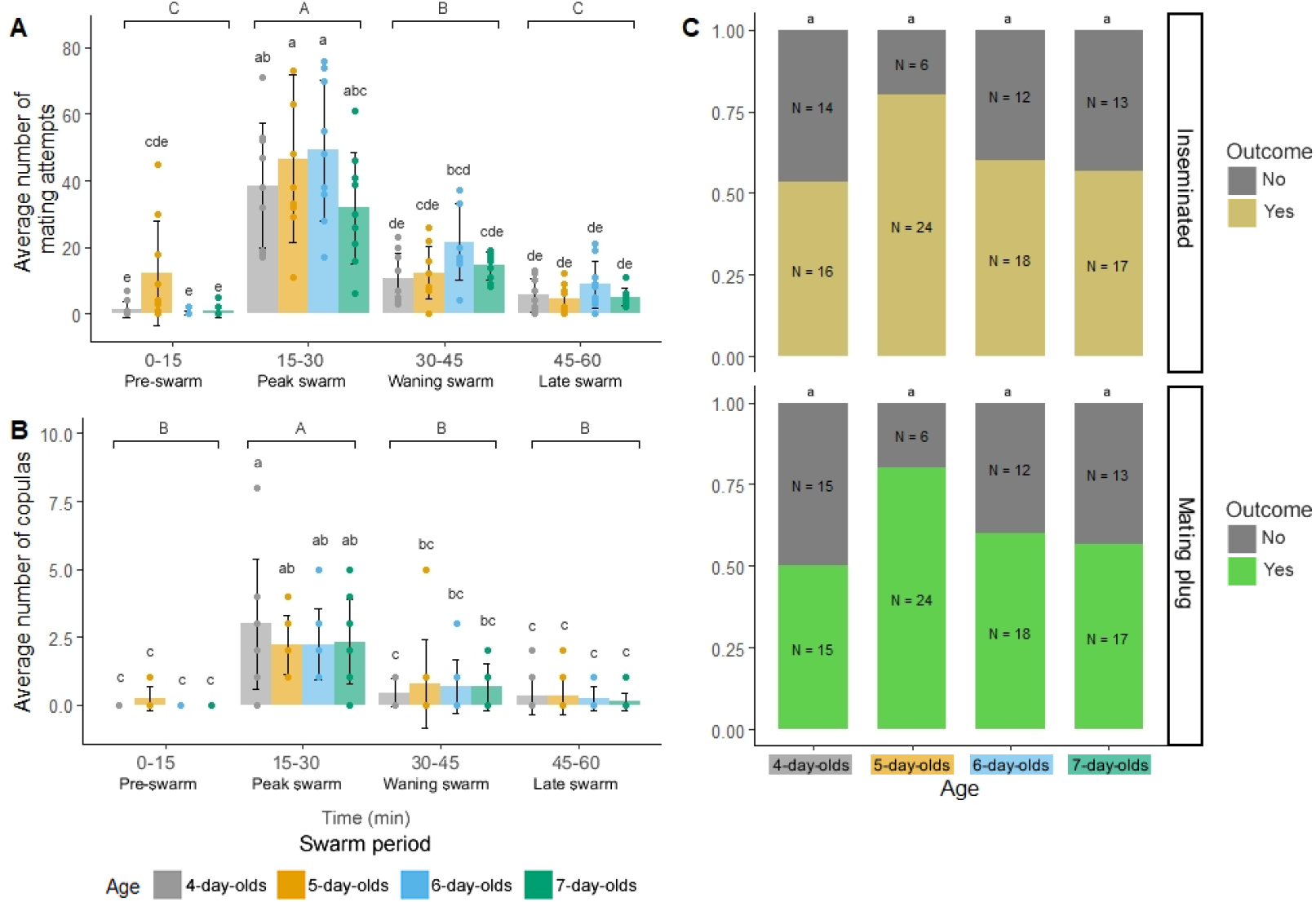
Mixed-sex swarm mating attempts, copulas, and mating outcomes vary by swarm period and age. Average number of mating attempts (**A**) and copulas (**B**) presented by swarm period and age. (**C**) The proportion of females that were inseminated (top graph) or that contained a mating plug (bottom graph) at the end of the swarm sequence presented by age. Bar plots in (**A**) and (**B**) display average ± SD values and letters above the bars represent Tukey pos-hoc test p-values for comparisons between swarm periods (uppercase) and ages (lowercase). Letters above the stacked bars in (**C**) represent Fisher’s exact test p-values for comparisons between ages.

Dissection of female spermathecae and transferred (i.e., internal) male mating plugs after swarming revealed that 75 out of 120 females (62.5%) were inseminated (range by age ≈50– 80%), with all but one also containing a mating plug (n=74; Fig. 4C; Supplementary Fig. S3). In nature, *An. gambiae* males achieve peak mating success at approximately 4–7 days post-eclosion at equivalent environmental conditions to those used in our experiments^53^. Despite a trend toward higher mating success in 5- and 6-day-old females compared to 4- and 7-day-olds, insemination (Fisher’s exact test, p=0.138) and mating plug (p=0.090) transfer rates were similar between ages. Successful matings (i.e., insemination occurred, n=75) were far lower than the total number of mating attempts (n=2,371), suggesting low mating success rates overall in our setup. Although most mated females were monandrous (i.e., singly mated, as evidenced by only one mating plug; n=65), a small minority were polyandrous (i.e., multiply mated, as evidenced by more than one mating plug; n=9; Supplementary Fig. S4). Taken together, these data show that swarm mating activity trends strongly correlate with swarm tone modulation patterns.

### Harmonic frequency differences of less than 50 Hz between male and female swarm tone harmonics occur frequently throughout mating interactions

Courting male-female mosquito pairs harmonize their wingbeat frequencies in a phenomenon known as harmonic convergence^27^. To examine this acoustic behavior during swarming, we tested for harmonic convergence (i.e., <5 Hz frequency differences) and harmonic frequency differences of less than 50 Hz (i.e., <50, <20, and <10 Hz) both in the early and late phases of acoustic mating interactions (Fig. 5; Supplementary Fig. S4). We detected harmonic convergence (n=35/240, 15%) or harmonic frequency differences less than 50 Hz (115/240, 48%) in 150 out of 240 (62.5%) tested interaction phases (Fig. 5, middle gray bars). These harmonic frequency difference outcomes occurred equally often during the early (Early interaction: Yes, n=77; No, n=43) and late mating interactions phases (Late interaction: Yes, n=73; No, n=47; Fig. 5; Fisher’s exact test, p=0.689), suggesting that harmonic frequency differences of less than 50 Hz commonly occur throughout acoustic mating interactions.

**Figure 5.**
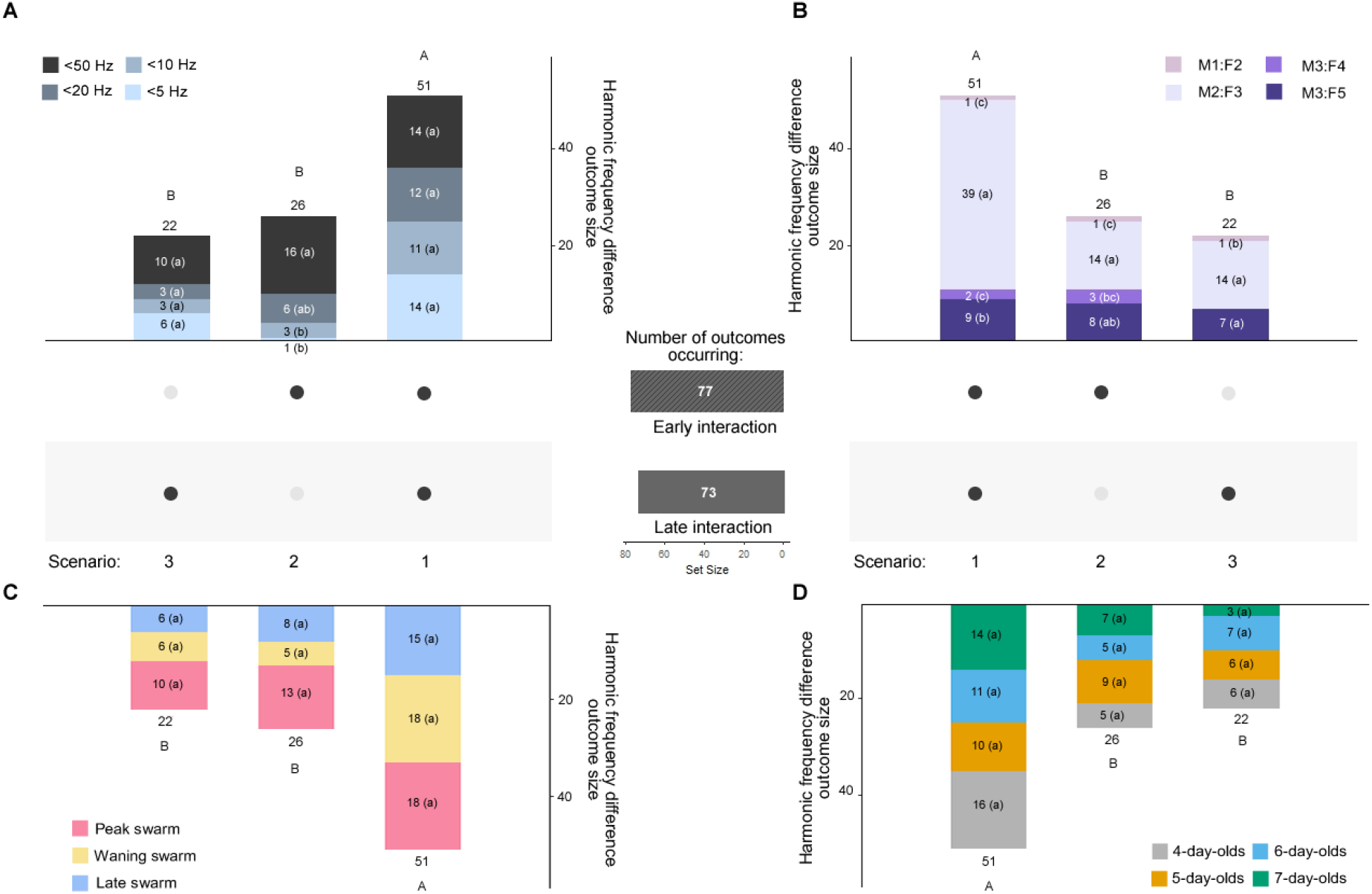
Harmonic frequency differences of less than 50 Hz between male and female flight tones occur throughout mating interactions. Harmonic frequency difference outcome scenarios (1–3, from most to least common scenario) analyzed by difference category (**A**), harmonic ratio (**B**), swarm period (**C**), and mosquito age (**D**). Middle gray bars represent the set size of frequency difference outcome scenarios by early or late interaction phase. For an illustration of the male and female flight tone sampling method during these interaction phases, see Supplemental Fig. S4. Letters above the stacked bars represent significant B-H post-hoc test differences between outcome scenarios (uppercase) and letters in parenthesis within bar categories represent significant B-H post-hoc test differences between frequency difference, harmonic ratio, swarm period, or age categories within outcome scenarios (lowercase). Abbreviations: M1, male fundamental flight tone harmonic; M2, male second harmonic; M3, male third harmonic; F2, female second harmonic; F3, female third harmonic; F4, female fourth harmonic; F5, female fifth harmonic.

In our analysis, harmonic frequency difference outcomes occurred in three possible scenarios: an outcome occurred both early and late in an interaction (scenario 1), only early in an interaction (scenario 2), or only late in an interaction (scenario 3). Frequency difference outcome scenarios differed significantly in their likelihood of occurrence (Fig. 5; Supplementary Table S7; Chi-square goodness-of-fit test= 20.067, p<0.001; B-H post-hoc test). In the most common scenario, males and females converged (<5 Hz) or displayed harmonic frequency differences of less than 50 Hz (<50, <20, or <10 Hz) both early and late in an interaction (scenario 1: n=51/120, 42.5%). Males and females attained these frequency differences only early in an interaction (scenario 2: n=26/120, 22%) as often as they did only late in an interaction (scenario 3: n=22/120, 18%). In the least common scenario (>50 Hz; Supplementary Data S1), mosquitoes did not achieve a harmonic frequency difference of less than 50 Hz in either mating interaction phase (n=21/120, 17.5%).

When less than 50 Hz frequency differences were obtained, they most often occurred at the M2:F3 harmonic ratio (Fig. 5B; Chi-square goodness-of-fit test= 22.727, 15.538, and 75.039 for scenarios 1, 2, and 3, respectively, p<0.001 for all tests; B-H post-hoc test). Frequency difference outcomes did not vary by swarm period or age (Fig. 5C, D; Chi-square goodness-of-fit test, p>0.05 for all tests). These results suggest that harmonic convergence and harmonic frequency differences of less than 50 Hz occur frequently through the entirety of acoustic mating interactions.

### Males attempt to mate in response to female baseline swarm tones, but not female mating interaction tones

Artificial flight tone stimuli have been used to examine acoustic mating behavior in mosquitoes^11, 26, 29, 30, 35, 39, 41, 62, 66^ as well as in midges^75^. To further dissect acoustic mating interactions during swarming, we tested swarming male flight responses to artificial female swarm tone audio playback emitted from a speaker (Fig. 6; Supplementary Fig. S2). Artificial female tones simulated alternating 5 sec playback periods of either female baseline swarm (550 Hz) or mating interaction (600 Hz) flight tone frequencies, which were derived from sampling single- and mixed-sex swarms, respectively (Supplementary File S1). Female audio playback experiments revealed that increases in male flight tone frequencies and amplitudes that occurred during swarming were especially pronounced during female baseline swarm tone playback periods (Fig. 6A, B; Supplementary Table S8 and S9; two-way repeated measures ANOVA, frequency: F(1:261)=23.6, amplitude: F(1:261)=24.36, p<0.0001 for both tests; Tukey post-hoc test). However in analogous experiments on females featuring playback of alternating artificial male tones (baseline swarm: 850 Hz; mating interaction: 900 Hz), female flight tone frequencies did not differ between male playback tone periods (Supplementary Fig. S5; Supplementary Table S10 and S11; two-way repeated measures ANOVA, F(1:254)=1.699, p=0.193; Tukey post-hoc test). We suspected that the increased male flight tone parameters during female baseline swarm tone playback periods resulted from mating interactions with the speaker, which were not induced in females in response to artificial male tones. We found that male flight tone frequencies (903.47 ± 38.79 Hz) and amplitudes (124.40 ± 7.29 dB) during interactions with the speaker (n=34) were higher than frequencies (843.01 ± 33.97 Hz) and amplitudes (100.49 ± 9.31 dB) during baseline swarming (n=34; Fig. 6C, D; paired t-tests, p<0.0001 for both tests).

**Figure 6.**
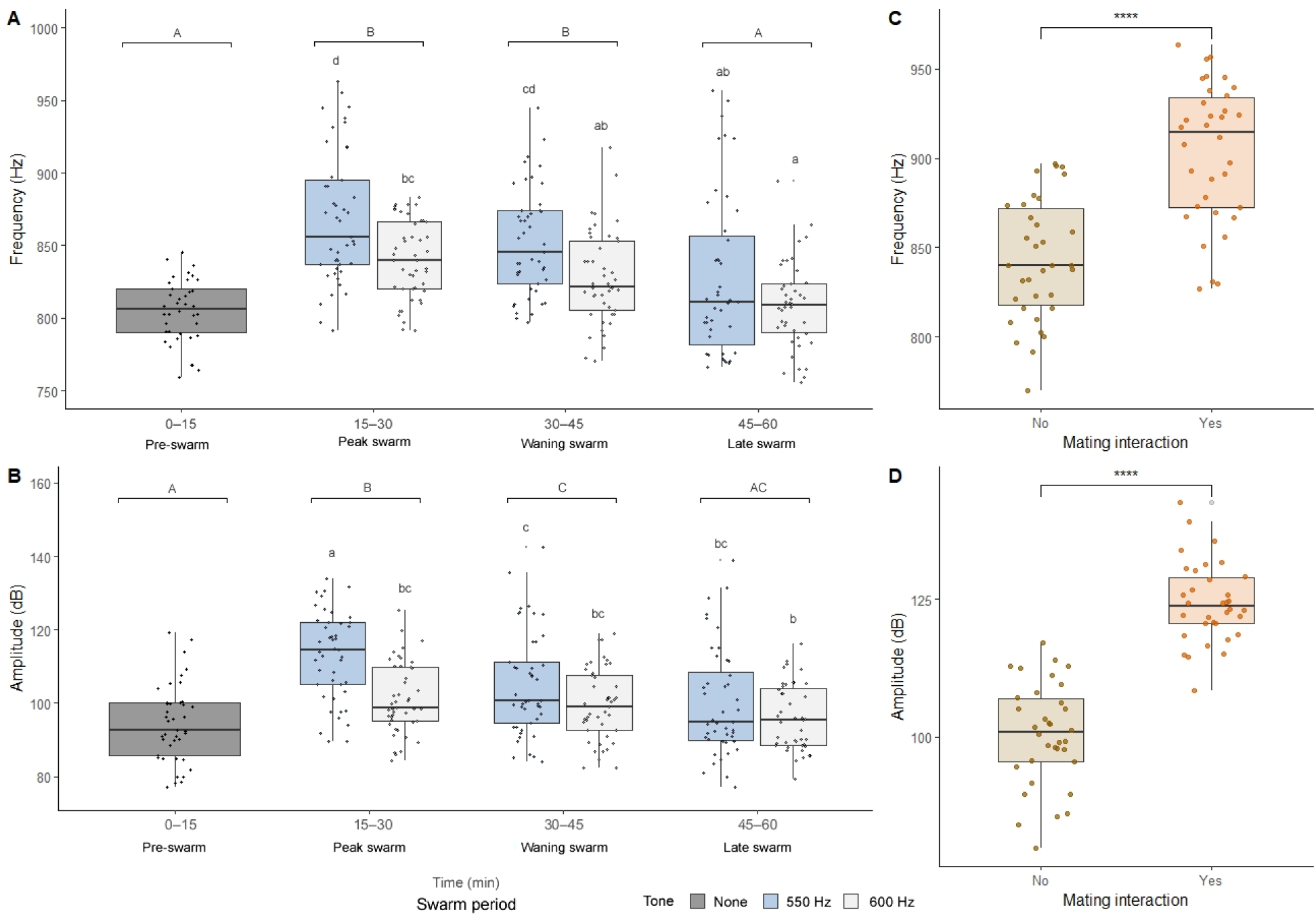
Males modulate flight tones in response to artificial female baseline swarm tones. Male flight tone frequencies (**A**) and amplitudes (**B**) by swarm period and artificial female flight tone playback period. Male flight tone frequencies (**C**) and amplitudes (**D**) in the presence or absence of an acoustic mating interaction. Letters above box and whisker plots represent significant Tukey pos-hoc test differences between swarm periods (uppercase) and female baseline (550 Hz) and mating interaction (600 Hz) audio playback periods (lowercase letters).

We next assayed for the presence or absence of male mating interactions at all 540 artificial female playback periods (270 baseline swarm tone periods and 270 mating interaction tone periods over three trials). In total, males engaged in acoustic mating interactions with the speaker in 163 of the 1620 playback periods (10%; Fig. 7A, B). Notably, all but one of these 163 interactions occurred during female baseline swarm tone periods, representing one in five (162/810, 20%) baseline tone periods (Fig. 7B; Chi-square goodness-of-fit test=174.63, p<0.0001). Consistent with mixed-sex swarm mating attempt patterns, male acoustic mating interactions climaxed during the peak-swarm period (11.56 ± 5.00) and declined through the waning (5.33 ± 3.46) and late (1.83 ± 0.98) swarm periods (Fig. 7C; one-way repeated measures ANOVA, F(2,21)=12.936, p<0.001; Tukey post-hoc test). Additionally, male-speaker interactions that met one, but not all three of our mating interaction criteria occurred overwhelmingly during female baseline swarm tone periods (Fig. 7D; Fisher’s exact test, p<0.0001 for each criterion). Together, these data show that male flight tone parameters increase due to swarming and acoustic mating interactions during swarming, which are only initiated in response to female baseline swarm tones. Furthermore, because females do not respond to playback of male flight tones alone, increases in female flight tones during swarming (Figs. 2, 3; Supplementary Fig. S4; Supplementary Table S2 and S3) appear to stem from other aspects of male presence, possibly including physical contact.

**Figure 7.**
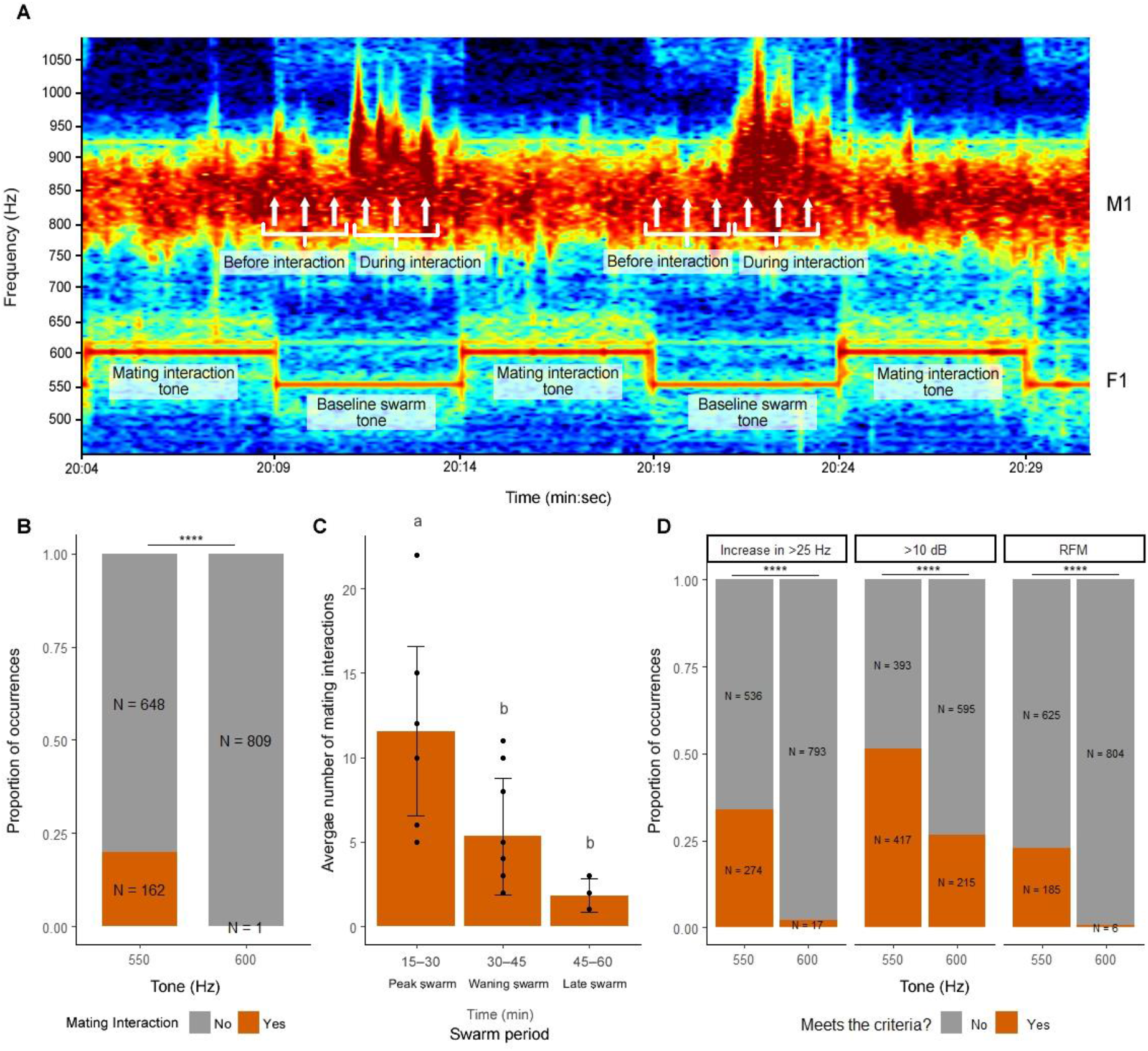
Male acoustic mating interactions occur in response to female baseline swarm tones, but not female mating interaction tones. (**A**) Spectrogram showing two instances of male acoustic mating interactions during female baseline swarm tone (550 Hz) playback periods and no instances of mating interactions during female mating interaction tone (600 Hz) playback periods. For harmonic frequency data in Fig. 8, male flight tone frequencies were sampled at three time points (white arrows) before and during each mating interaction. (**B**) Proportion of male acoustic mating interactions by female artificial swarm tone playback period. (**C**) Average number of male acoustic mating interactions by swarm period. (**D**) Proportion of male flight tone samples that met each of the three male mating interaction criteria by female playback period. For Chi-square goodness-of-fit test in **B** and Fisher’s exact test in **D**: **** p<0.0001. In **C,** bar plots in display average ± SD and letters above bars represent Tukey post-hoc test p-values for comparisons between swarm periods. Abbreviations: M1, male fundamental flight tone harmonic; F1, female fundamental flight tone harmonic.

### Harmonic frequency differences of less than 50 Hz occur frequently between male and artificial female baseline swarm tones and mating interaction tones

To gain additional insight into harmonic frequency differences during swarming, we tested for harmonic convergence (<5 Hz) and harmonic frequency differences of less than 50 Hz (<50, <20, and <10 Hz) at the M2:F3 harmonics between male and artificial female baseline swarm tones and mating interaction tones (550 Hz and 600 Hz artificial female baseline and interaction tones, respectively; Fig. 8). We detected harmonic convergence or frequency differences of less than 50 Hz either before or during all male-speaker acoustic mating interactions (n=163/163, 100%). Immediately before mating interactions (Fig. 8, striped horizontal bars), harmonic frequency difference outcomes were almost twice as likely to occur between male baseline swarm tones and artificial female baseline swarm tones (n=136/163, 83.44%) compared to female mating interaction tones (n=79/163, 48.47%). Likewise, during mating interactions (Fig. 8, solid horizontal bars), convergence was nearly three times more likely to occur between male interaction tones and female mating interaction tones (n=146/163, 89.57%) compared to female baseline swarm tones (n=57/163, 34.97%). We therefore detected a strong association between harmonic difference outcomes and female baseline swarm tones before mating interactions and female mating interaction tones during mating interactions (Fisher’s exact test, p<0.0001).

**Figure 8.**
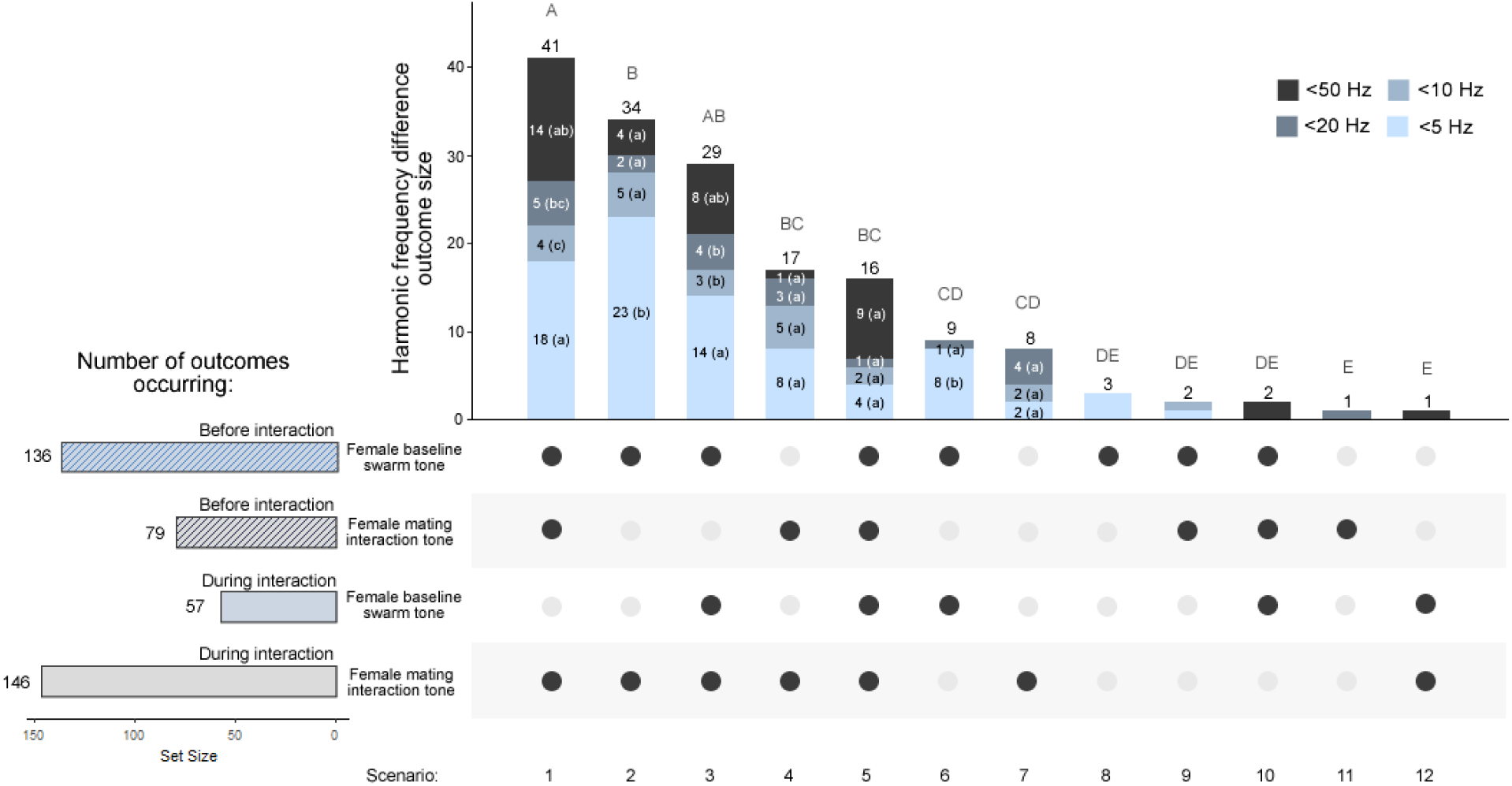
Harmonic frequency differences of less than 50 Hz between male and artificial female flight tones occur before and during mating interactions. Harmonic frequency difference outcome scenarios (1–12, from most to least common scenario) were analyzed by difference category (vertical stacked bars). Horizontal bars represent the set size of frequency difference outcome scenarios by interaction phase: immediately before (striped bars) or during (solid bars) acoustic mating interactions, during either the female baseline swarm tone (blue bars) or mating interaction tone (gray bars) playback periods. For an illustration of the male flight tone sampling method during these interaction phases, see Fig. 7A. Letters above the vertical stacked bars represent significant B-H pos-hoc test differences between outcome scenarios (uppercase) and letters in parenthesis within bar categories represent significant B-H post-hoc test differences between frequency difference categories within outcome scenarios (lowercase).

Harmonic frequency difference outcomes for interacting males fell into one of twelve outcome scenarios, which differed significantly in their likelihood of occurrence (Fig. 8; Chi-square goodness-of-fit test= 159.97, p<0.0001; B-H post-hoc test). Notably, approximately three-quarters of all scenarios (120/163, 74%) involved two outcomes (see scenarios 1–3 and 5): 1. males converging or reaching frequency differences of less than 50 Hz with both the female baseline swarm tone prior to the interaction; and 2. males converging or reaching frequency differences of less than 50 Hz with the female mating interaction tone during the interaction. Indeed, harmonic convergence (<5 Hz) was typically the most common difference category in scenarios that involved these two outcomes (Chi-square goodness-of-fit test=13.73 and 33.53 for the first and second outcome, respectively, p<0.01 for both tests; B-H post-hoc test). Together with mixed-sex swarm data, these results suggest that males and females swarm at or near harmonic convergence and adjust their flight tones during mating interactions to maintain optimal frequency differences at higher frequencies.

## Discussion

Acoustic communication is essential for mating in mosquitoes^26, 31, 41, 76, 77^ as well as other disease vectors^78^ and insects^74^. Here, we provide the first detailed analysis of *An. gambiae* flight tone frequencies and harmonic convergence in free-flying individuals across a complete swarm sequence, coupling these acoustic phenomena to their underlying mating behaviors. We show that both males and females dynamically modulate their flight tones during swarming, particularly when engaging in mating interactions. We also provide evidence indicating harmonic convergence coordinates swarming flight and individual mating interactions to optimize detection of females and acoustically mask mating pairs.

### Harmonic convergence and coordination of mosquito swarm mating activity

This study is the first to examine harmonic convergence in free-flying male and female mosquitoes in mixed-sex swarms and in single-sex swarms with artificial flight tone playbacks across an entire swarm sequence (Fig. 5–8). By assaying for harmonic frequency differences between male and female tones both immediately prior to and during mating interactions, rather than only before^10, 39^ or during^40, 67^ interactions, we found that overtone differences of less than 50 Hz occur as often between collective male and female baseline swarm tones as they do between individual male and female mating interaction tones. We propose that males and females swarm near harmonic convergence at the male second and female third harmonic overtones (M2:F3; Fig. 2A) to enable efficient sex recognition^28^. These harmonics are not only among the most commonly reported converging overtones in mosquitoes^10, 26, 28, 30, 31, 39, 47, 49^ but are also those for which *An. gambiae* male and female antennae are particularly sensitive^31, 39^. The clustering of male flight tone frequencies observed here during swarming (Figs. 1 and 2; Supplementary Table S2 and S3) are consistent with the suggestion by others that males tightly coordinate their swarming flight^4, 30, 47, 79^ and collectively synchronize their flight tones to minimize acoustic interference and efficiently detect females^30, 66, 75, 79^.

In our audio playback experiments, swarming males (≈850 Hz) responded vigorously to playback of female swarm tones (550 Hz, ≈300 Hz difference), and yet a frequency difference of only ≈50 Hz was sufficient to render female mating interaction tones (600 Hz, ≈250 Hz difference) almost completely unattractive to males (Figs. 6 and 7). Numerous studies suggest that males tune into the frequency differences between male and female fundamental flight tones, also known as acoustic distortion products, rather than listening to female tones themselves^10, 28, 31, 66^. Thus, by adjusting their flight tones during mating attempts (≈900 Hz) to maintain the same frequency difference with interacting females (600 Hz, ≈300 Hz difference), males could still detect females during a mating attempt despite their increased flight tone frequencies. Swarm activity coordination based on frequency difference-mediated hearing could enable males to rapidly detect, locate, and even amplify the swarm tones of available females while avoiding mating attempts with females engaged in courtship flight with other males^28, 29, 31, 33, 34^. Thus, in mating systems with highly skewed sex ratios in which many males compete for few females^3^, we propose that higher frequencies during mating interactions serves to screen out courting male-female pairs from other nearby male suitors. This mating coordination strategy is likely conserved across mosquitoes, as increases in flight tone frequencies during mating interactions were also recently observed in *An. gambiae* and *An. coluzzii*^39^, *An. albimanus*^47^, *An. darlingi*^48^, and *Culex quinquefasciatus*^10, 66^, and increases in flight speed in response to artificial tone playback were observed in another study in *An. gambiae* males and *Ae. aegypti* males and females^35^.

Male hearing can be highly entrained to female frequency differences during swarming^31, 34^. We hypothesize that swarming males are unable to hear distortion products of males and females during a mating interaction, and hence are unresponsive to interacting female tones^31, 66^. This is supported by the fact that females swarm at ≈550 Hz, which is just below the uppermost frequency boundary of male mating behavior thresholds in *An. gambiae*^39^ and other mosquitoes^10, 39, 80^. This would also mean that female mating interaction tones of ≥600 Hz would be just beyond the male behavior threshold boundary. Mechanistically, we propose that by courting at tones that are only 50 Hz higher than swarm tones, female tones during mating interactions function as “masking” tones that suppress male mating attempts by interfering with the male’s ability to detect female baseline swarm (i.e. “probe”) tones^66^. This phenomenon is especially effective when the probe and masking tones are close in frequency^66, 81^, which was the case in our artificial female tone playback experiments. Interacting mating pairs in our study were also louder (typically ≥10 dB) than their swarming counterparts, and increased flight tone amplitudes elicit stronger phonotactic responses in males^35^. Increased flight tone amplitudes during mating interactions may therefore serve as an additional signal by which males may discern between interacting females and available females^66^.

Consistent with previous studies in free-flying *An. gambiae*^35, 62^, and in contrast to behavior in *Ae. aegypti*^30, 35^, our data show that females do not initiate mating interactions in response to male flight tones alone (Fig. S5). Previous studies in *An. gambiae*^26^ and *Ae. aegypti*^30^ documented harmonic convergence responses in both males and females when presented with live individuals and artificial flight tones of the same or opposite sex. However, this apparent discrepancy is almost certainly due to differences between free-flying and tethered mosquito analyses, as tethering permits detection of active flight adjustments over longer periods of time (e.g., ≈10–60 s^26, 30^) compared to relatively brief free-flying interactions (≈1–2 sec^10, 35, 39, 47^). Our data confirm observations of strong phonotactic responses in *An. gambiae* males to female flight tones^35^, particularly around 500 ± 50 Hz (Fig. 6)^4, 11, 31^. Together, these observations suggest mating interactions and accompanying acoustic phenomena like harmonic convergence are initiated by stereotypic male responses to female flight tones, such as rapid frequency modulation^10^, which in turn trigger stereotypic female responses to male contact at the point of male interception^10, 40^, such as “startle response” flight^29^. Our findings suggest a more expansive scope for harmonic convergence, which appears to depend upon swarming flight coordination^79^, flight tone coordination during stereotypic courtship behaviors^10^, and finally active male-female flight adjustments made during the latter phases of mating interactions^40^.

### Mating-dependent and -independent swarm acoustic modulations

We decoupled swarm acoustic modulations from their underlying mating behaviors by examining single-sex swarms (Fig. 1) and coupled observations of these same phenomena in mixed-sex swarms (Figs. 2–4). Collectively, we document a transient increase in flight tone parameters due to swarming and mating activity shortly after sunset that tapered off gradually over time. Our observed swarm latency, duration, and mating activity were remarkably similar to measurements obtained from *Anopheles* mosquitoes in field^4, 6, 7^ and laboratory studies^4, 11^. We also detected differences in flight tones and mating activity related to mosquito age. Our finding that flight tone frequencies tend to increase with age confirm earlier trends observed in *Ae. aegypti*^24^. Similar to previous reports in *An. gambiae*^11, 53^, we found that mating activity and mating success tended to peak at around five to six days post-eclosion. We posit that mosquito flight and mating activity are dependent in large part upon mosquito age and energy reserves at the time of swarming^52^. Future studies could explore these trends in swarm acoustics and mating further by utilizing a broader range of ages and variable adult nutrition.

The free-flying male and female flight tone frequencies observed here (Supplementary Table S2 and S3) were substantially higher compared to previous data derived from tethered mosquitoes^26^, but were consistent data from an earlier comparative study of free-flying *Anopheles* spp. flight acoustics^36^. Numerous studies have reported mosquito flight tones from single, pairs, or groups of tethered adults^26, 28–31, 45, 79, 82–84^, free-flying adults^24, 32, 35–37, 49, 66, 85^, or a combination of tethered and free-flying adults^10, 21, 25, 39–42, 47, 48, 62, 67, 80^. Fewer studies have examined mating activity in swarms^6, 86, 87^ or performed audio recordings of swarming individuals in free flight^10, 35, 36, 39, 47, 49, 62, 66, 85^, and none prior to the present study have done so simultaneously or over an entire swarm sequence. Overall, the findings presented in this study, along with other recent mosquito acoustics work^10, 39, 47, 48, 66^, underscores the importance of examining mosquito flight tones and mating activity in free-flying individuals, ideally during their swarming period.

### Implications for acoustic-mediated mosquito control methods

The data presented here reveal ample opportunities for acoustic manipulation of mosquito auditory systems and swarm behavior, a novel and rapidly developing dimension of vector and disease control efforts^16, 88, 89^. Because *Anopheles* species swarm relative to contrasting ground markers^4, 16, 53, 87, 90, 91^, often at that same location within and across seasons^6, 92^, swarms provide an attractive biological opportunity for targeted vector control^16, 31, 93, 94^. Mosquito swarm acoustics show promise for trapping and surveillance purposes^14–24^ and could be implemented alongside existing control methods involving sterile insect techniques or genetically-modified male mosquito releases^16, 91^. Our study provides reference data against which the mating behavior of modified lines can be compared^62, 82^. This study also furnishes an experimental template that could be adapted to create artificial flight tones that either stimulate or disrupt swarming in nature. For instance, artificial tones could be designed for use in push-pull acoustic systems that induce positive^35^, or negative^95^ phonotactic responses in individuals, depending on the application and desired outcome.

### Limitations and conclusions

Future research can significantly expand upon the biological insights into mosquito swarm acoustics presented here. For instance, our hypothesis that swarming males are unable to hear distortion products of males and females engaged in a mating interaction could be tested neurophysiologically by measuring responses of the Johnston’s Organ antennal nerve^80^. Future work could significantly expand upon our laboratory-based experiments by examining field-collected mosquitoes in field or semi-field environments^86, 90^, ideally with more spacious enclosures, larger swarms, and higher male:female sex ratios^3, 4, 6^ than those employed here. Finally, our study included manual analyses of swarm acoustics, which despite their precision, require extensive training and are time intensive. To increase data throughput, future analyses of swarm acoustics would benefit from automation^30^.

The data presented here provide compelling evidence that harmonic convergence during mosquito swarming serves a critical function in the coordination of emergent acoustic mating behavior^8, 79, 96^. This study lays the groundwork for future investigations into the nature and biological function of mosquito swarm acoustics in *An. gambiae* and other key disease vectors. The fine-scale tuning of acoustic mating responses described here could be leveraged to improve current vector surveillance and disease control methods as well as pave the way for new ones.

## Acknowledgements

This study was supported by a National Institute of Allergy and Infectious Diseases (www.niaid.nih.gov) grant R01AI095491 to LCH and Dr. Mariana F. Wolfner. SSGC received support from a Fogarty International Center (www.fic.nih.gov) Translational Research Development for Endemic Infectious Diseases of Amazonia grant 2D43TW007120-11A1 to Dr. Joseph M. Vinetz and Dr. Dionicia Gamboa Vilela. We acknowledge the Research Experience for Peruvian Undergraduates program for facilitating SSGC’s internship at Cornell. KSP received undergraduate research funding from a supplement to NIFA Hatch -NYC 139443. Finally, we are grateful to Dr. Itai Cohen and Dr. Samuel Whitehead of Cornell University for technical feedback and use of their insect flight recording cage for the swarm experiments.

---

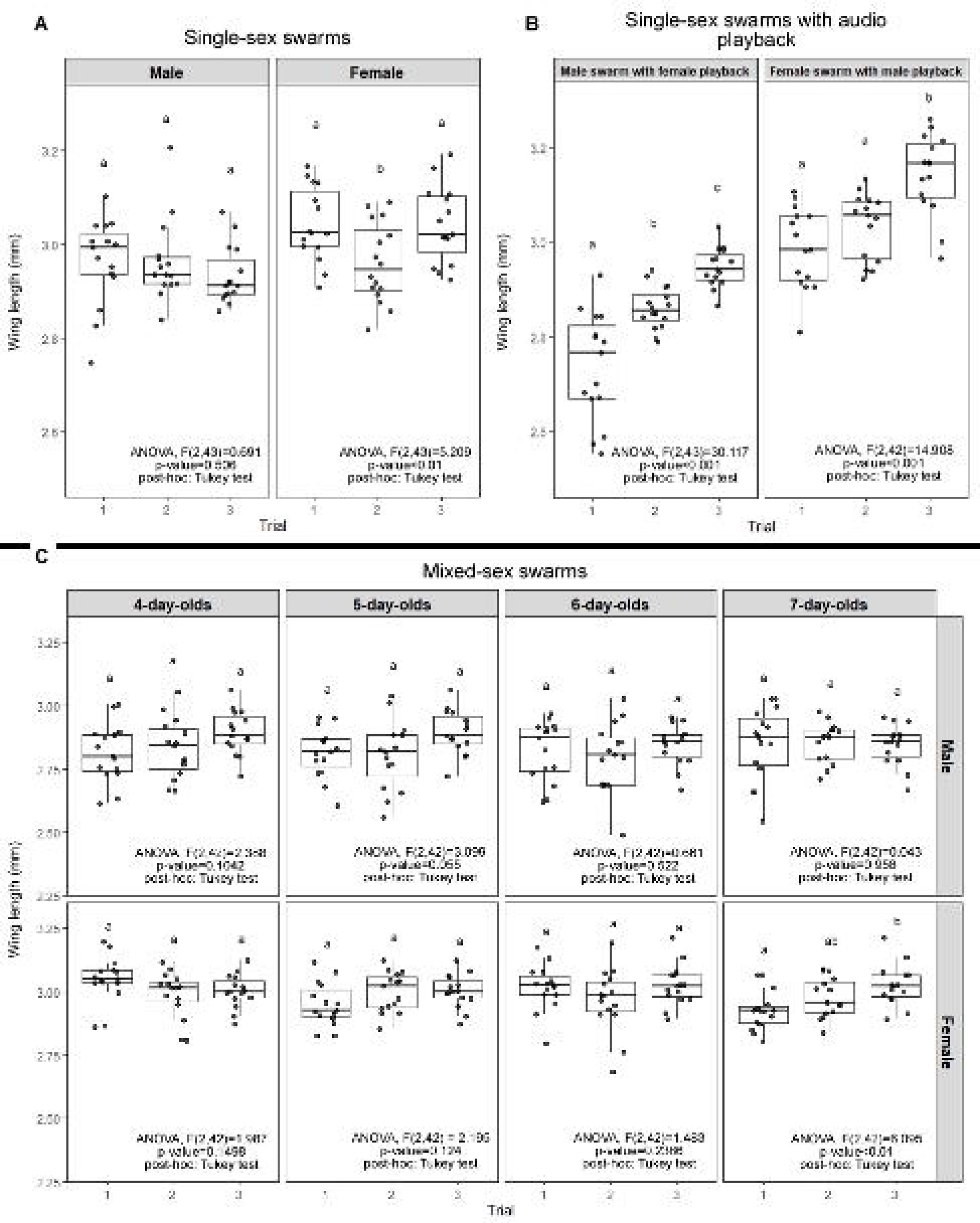

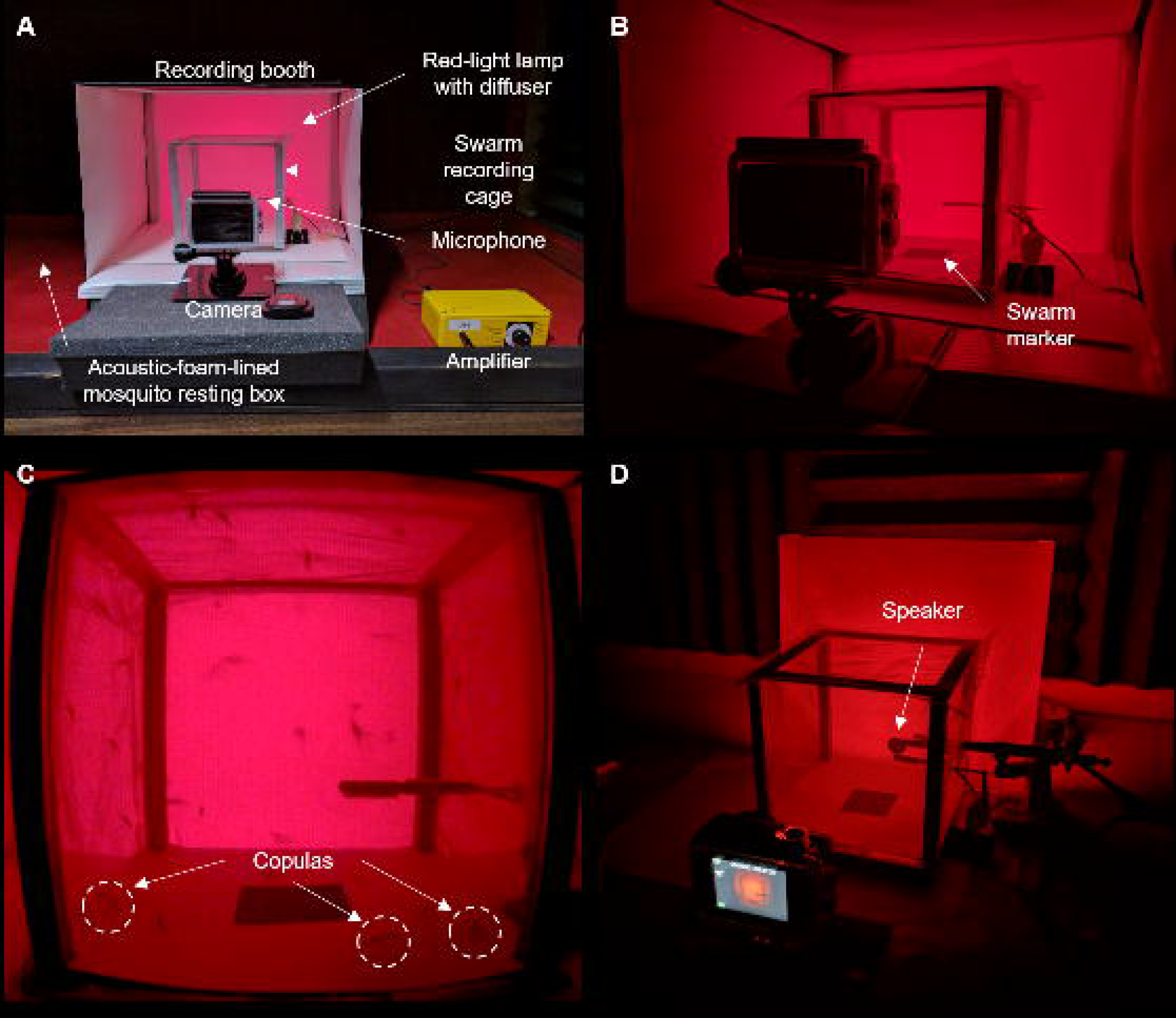

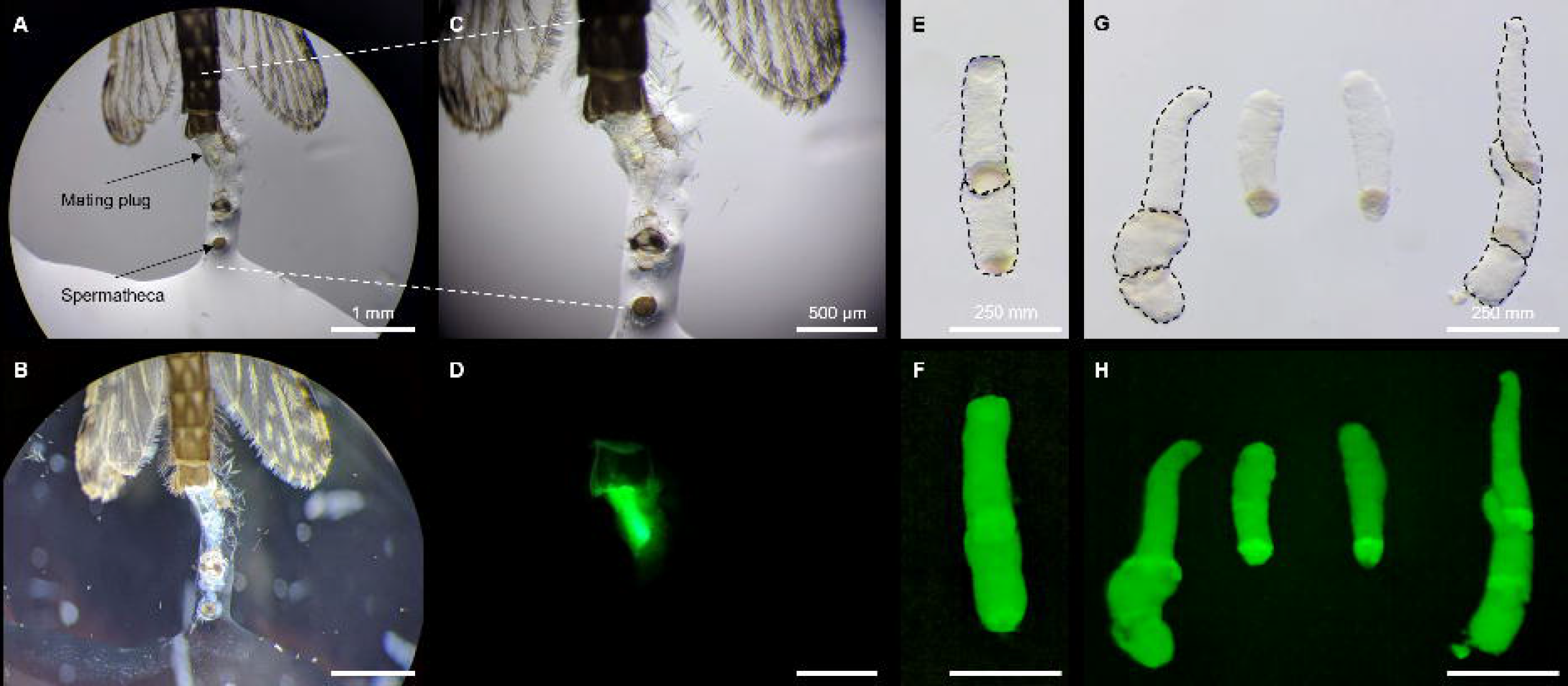

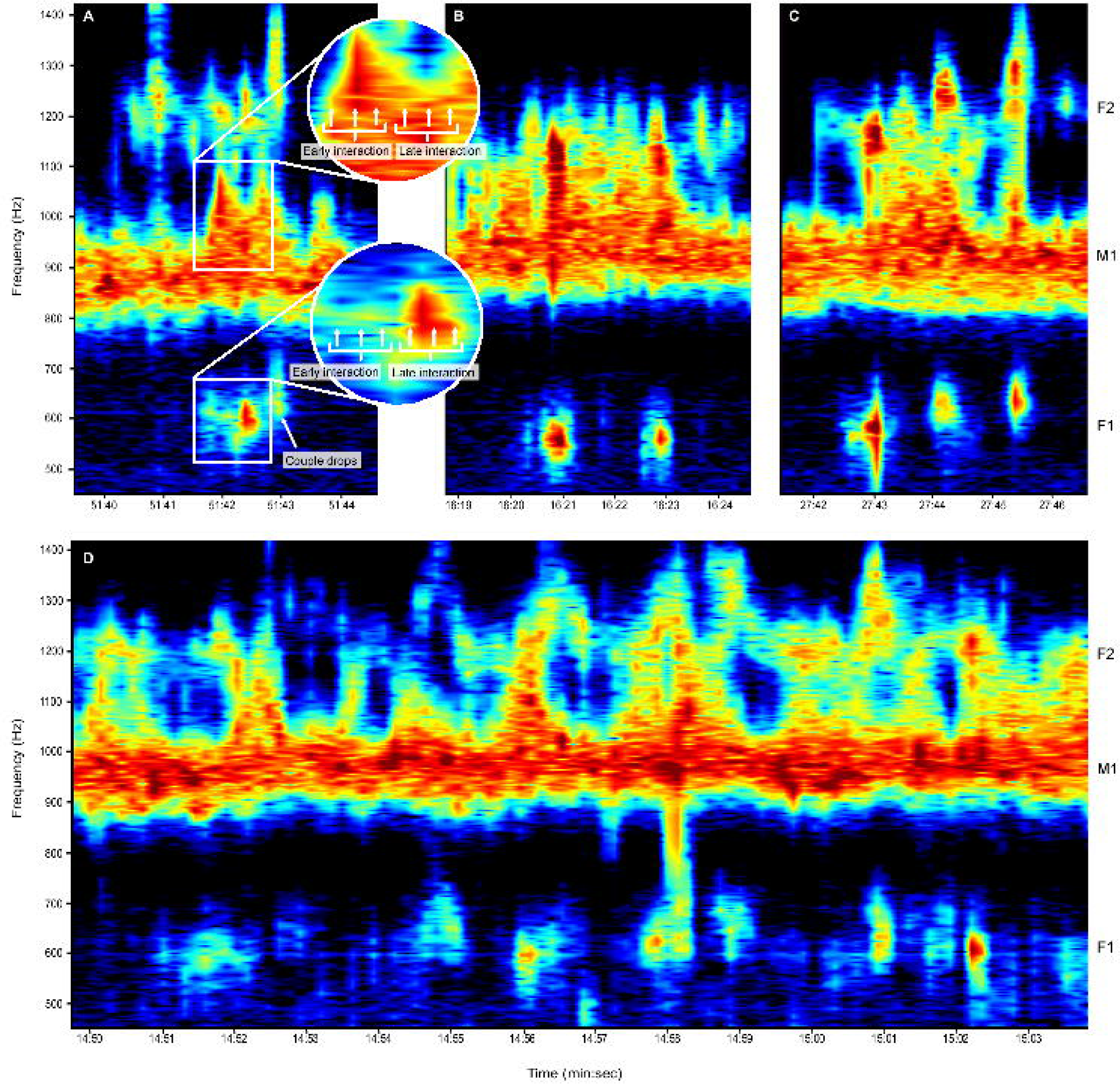

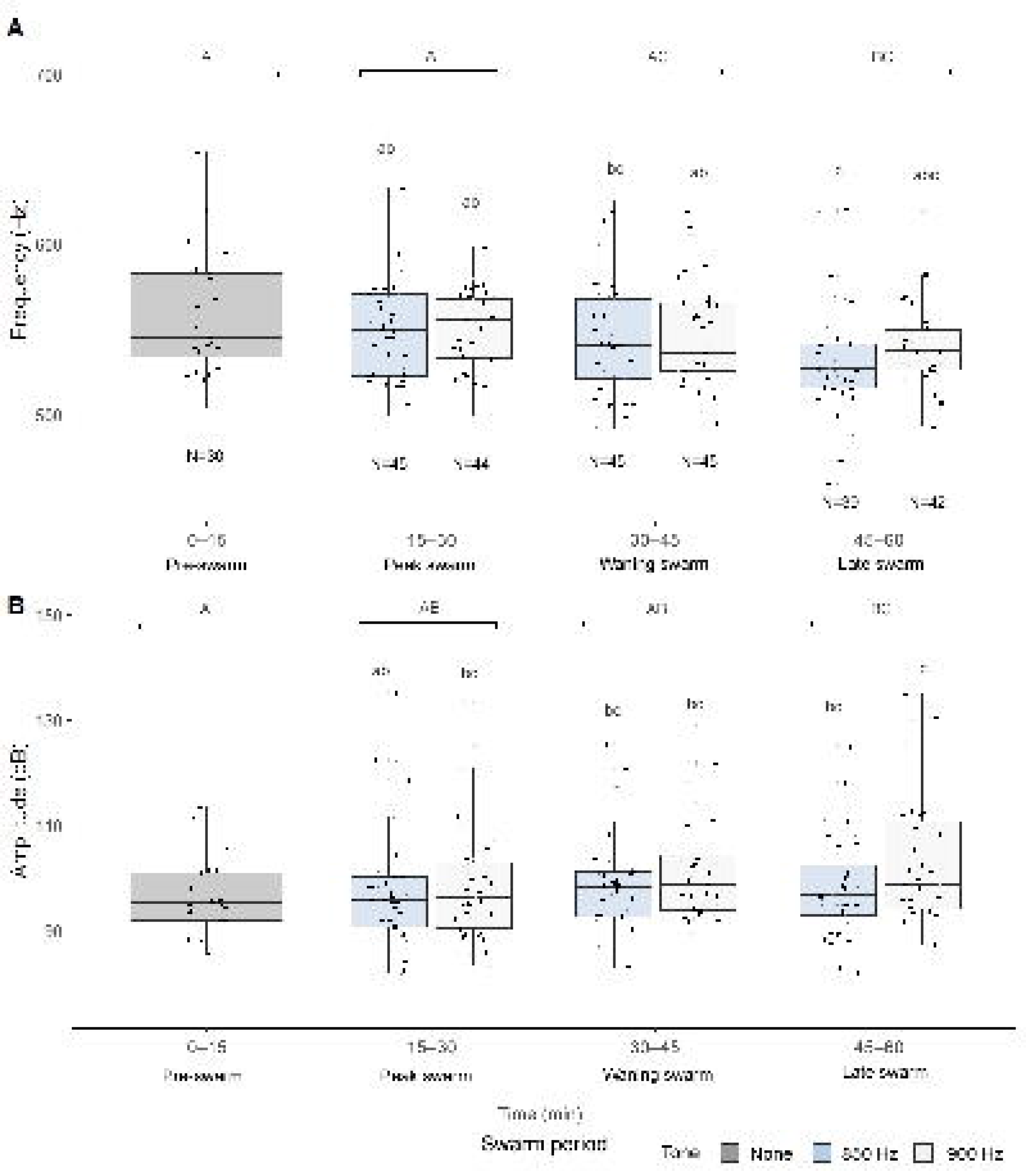

