## Supplementary Fig. for "Synchronized swarming: Harmonic convergence and acoustic mating dynamics in the malaria mosquito *Anopheles gambiae*"

**Supplementary figures**

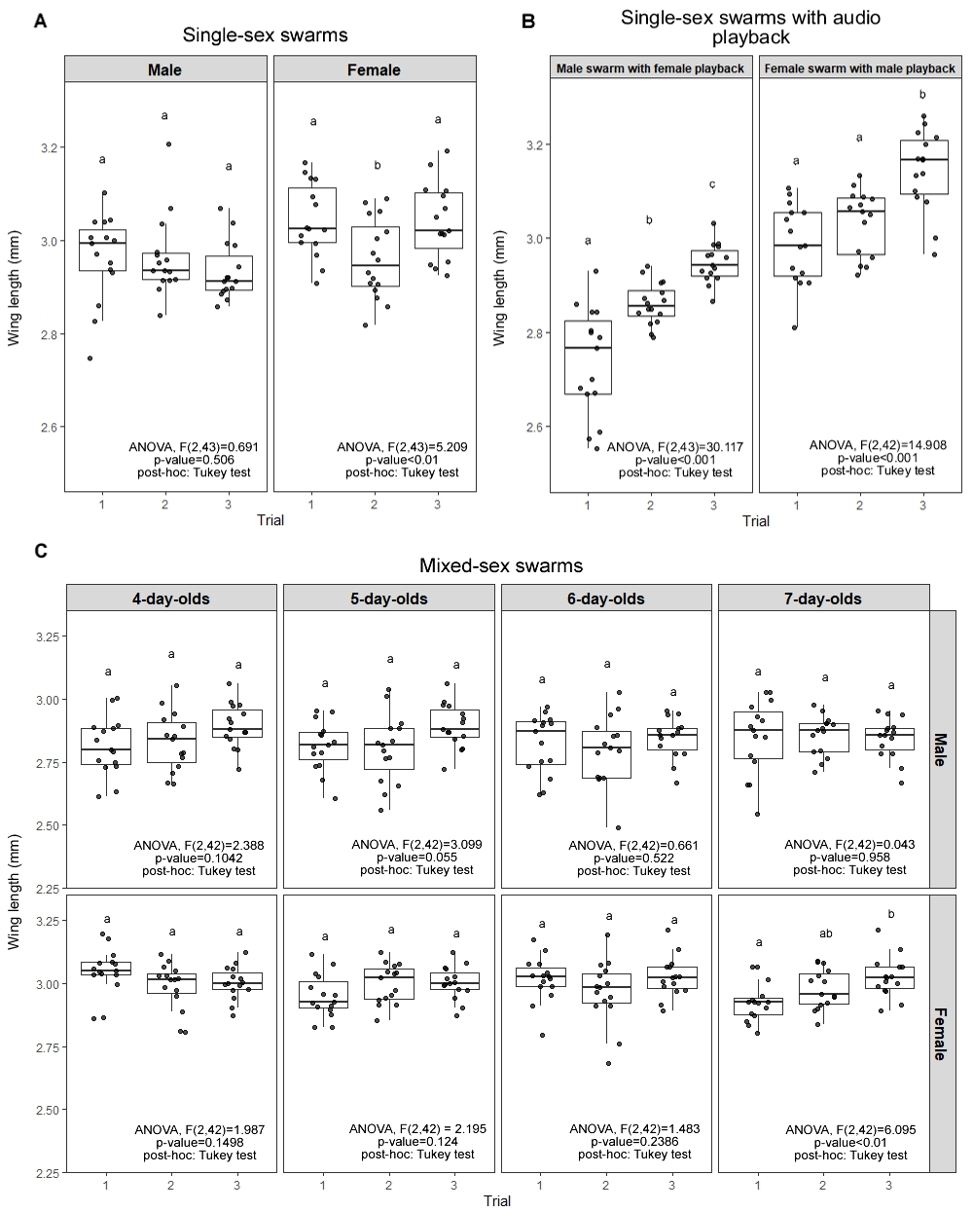

**Figure S1. Male and female wing lengths by experiment, trial, and/or age.** Wing lengths for single-sex swarm experiments (**A**), single-sex swarm experiments with audio playback (**B**), and mixed-sex swarm experiments (**C**). Lowercase letters above box and whisker plots denote Tukey post-hoc test p-values for comparisons between trials.

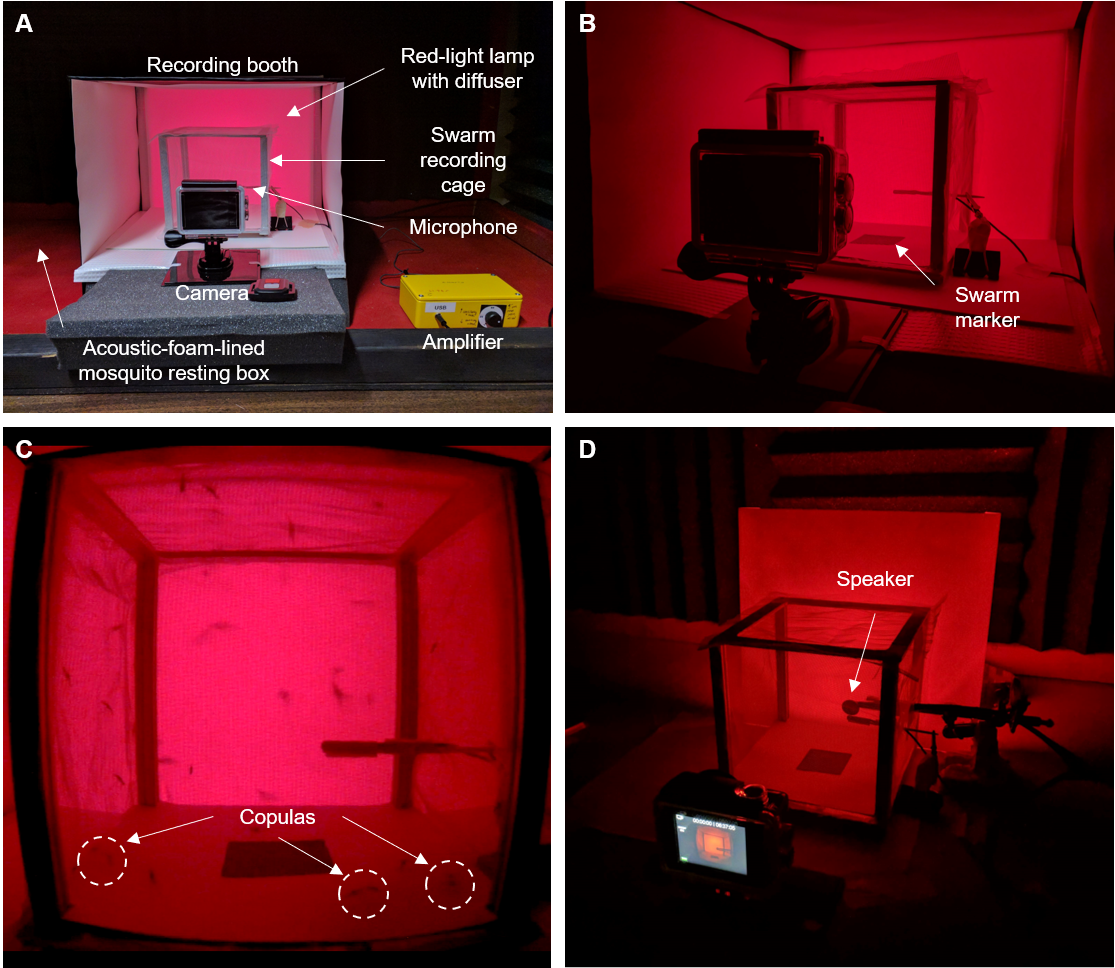

**Figure S2. Swarm recording setup.** (**A**) Swarm recording setup was housed in an acoustic-foam-lined mosquito resting box and included a cage with mosquito net walls and a red-light lamp with a diffuser inside of a small recording booth for lighting. A microphone attached to an amplifier recorded swarm audio and a camera recorded video. (**B**) The microphone was inserted through the cage net and over a swarm marker to optimize swarm audio capture. (**C**) Still image from a swarm video showing swarming mosquitoes and three mating male-female pairs in copula on the cage floor (dashed circles). (**D**) During audio playback experiments, a speaker placed 2 cm from the microphone emitted artificial male or female flight tones. Ambient lights are on only in (**A**) and red-light lamp is on in (**A**–**D**).

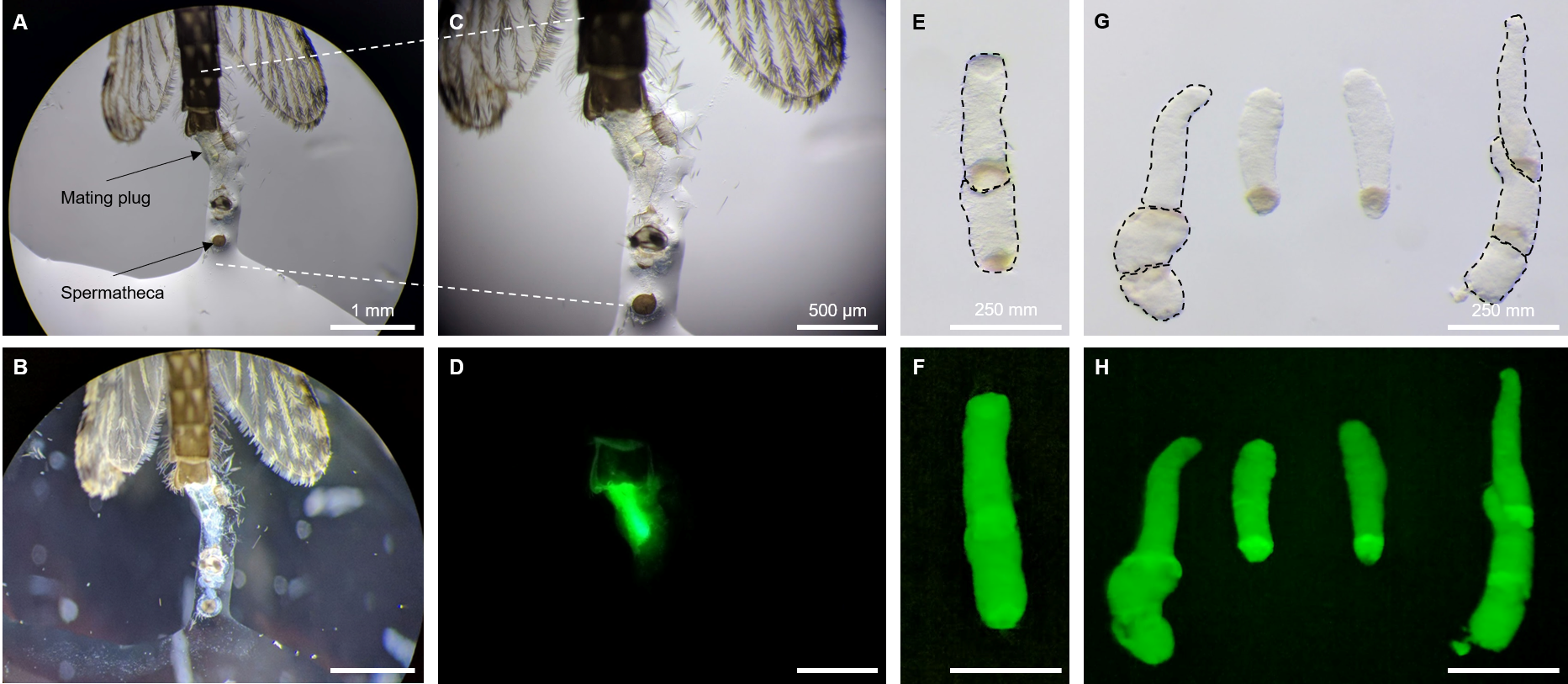

**Figure S3. Spermatheca and mating plug dissection following mixed-sex swarm recordings.** (**A**) Dissected female abdomen imaged under oblique illumination showing spermatheca and mating plug, which were used to determine female insemination and mating plug transfer status. (**B**) Dissection in panel (**A**), only imaged under darkfield illumination to reveal sperm inside the spermatheca. (**C**) Higher magnification of panel (**A**) dissection image. (**D**) Dissection in panel (**C**), only imaged under fluorescence Illumination to show mating plug autofluorescence. (**E**) Two mating plugs aligned on top of one another from a single dissection imaged under brightfield illumination, indicating a polyandrous (i.e., multiply mated) female. (**F**) Mating plugs in panel (**E**) showing autofluorescence. (**G**) Series of mating plugs from four dissections imaged under brightfield illumination, indicating both monandrous (i.e., singly mated) and polyandrous females. (**H**) Mating plugs in panel (**G**) showing autofluorescence.

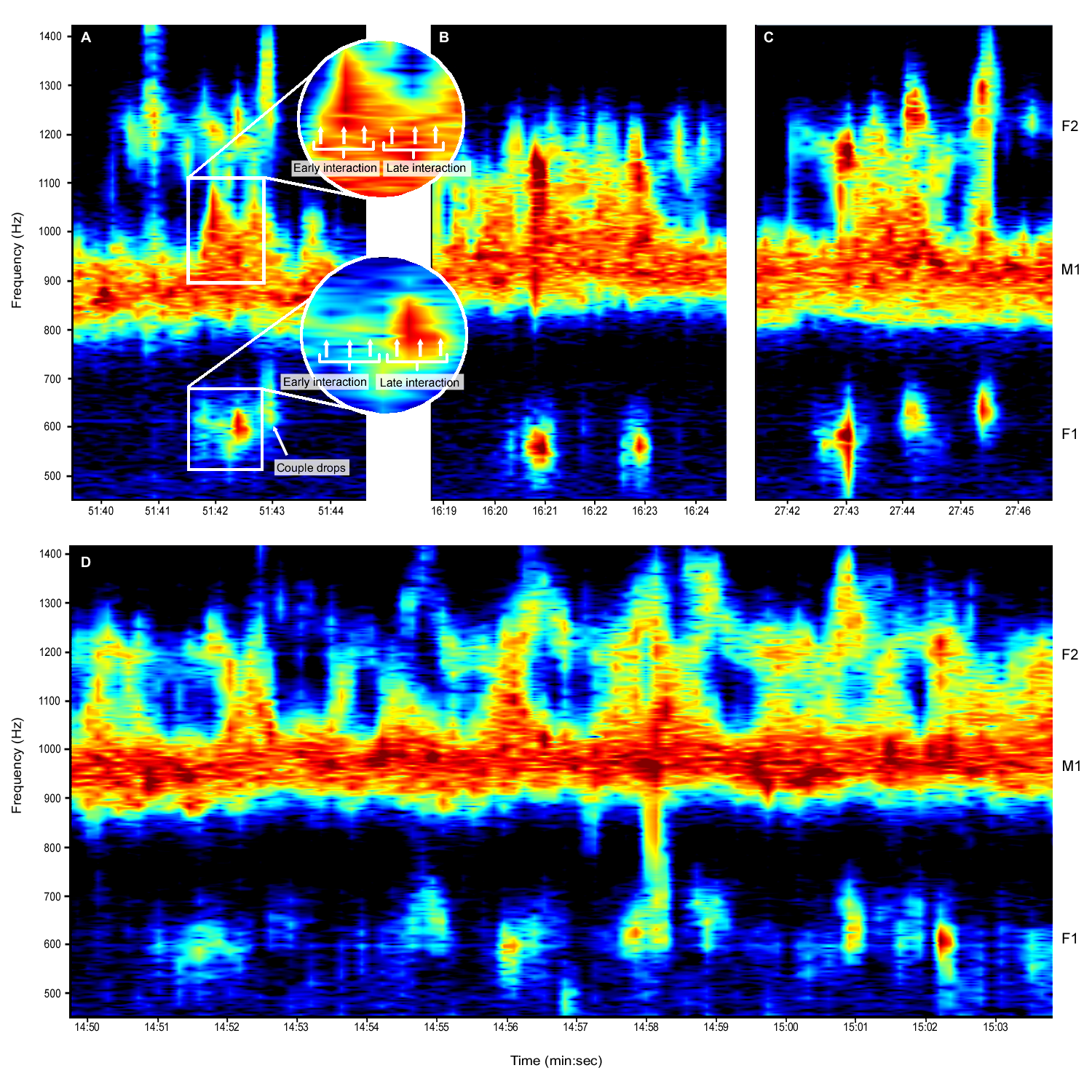

**Figure S4**. **Mixed-sex swarm acoustic mating interactions.** (**A**) Male (top box) and female (bottom box) acoustic mating interaction flight tone frequencies were sampled at three time points (white arrows) in the early and late interaction phases (zoomed in circle inserts). Mating interactions typically ended once male and female couples dropped out of the swarm. When acoustic mating interactions contained multiple segments (**B**, **C**) or occurred in a sequence of multiple interactions (**D**), only the first segment of the interaction closest to the sampling time point was analyzed. Abbreviations: M1, male fundamental flight tone harmonic; F1, female fundamental flight tone harmonic; F2, female second harmonic.

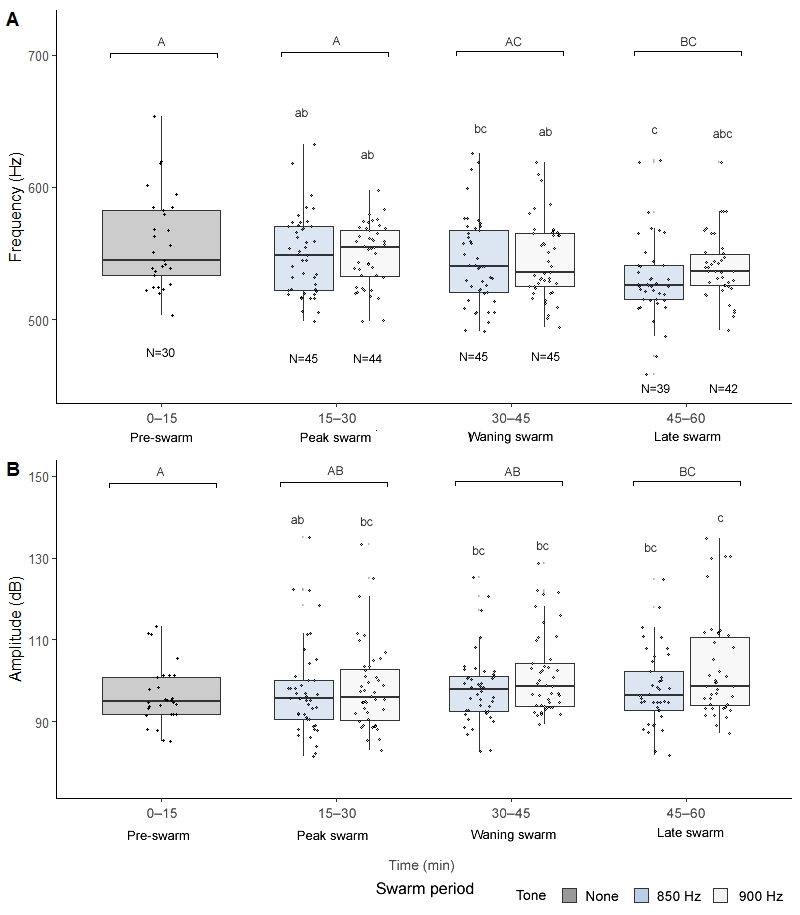

**Figure S5.** **Females do not modulate flight tones in response to artificial male flight tones.** Female flight tone frequencies (**A**) and amplitudes (**B**) by swarm period and artificial male flight tone playback period. Letters above box and whisker plots represent significant Tukey pos-hoc test differences between swarm periods (uppercase) and male baseline (850 Hz) and mating interaction (900 Hz) audio playback periods (lowercase letters).
